# Nucleus accumbens cholinergic interneurons oppose cue-motivated behavior

**DOI:** 10.1101/520817

**Authors:** Anne L. Collins, Tara J. Aitken, I-Wen Huang, Christine Shieh, Venuz Y. Greenfield, Harold G. Monbouquette, Sean B. Ostlund, Kate. M. Wassum

**Affiliations:** Dept. of Psychology, UCLA, Los Angeles, CA 90095; Dept. of Chemical Engineering, UCLA, Los Angeles, CA 90095, USA; Dept. of Anesthesiology and Perioperative Care, UCI, Irvine, CA 92697, USA; Brain Research Institute, UCLA, Los Angeles, CA 90095, USA

**Keywords:** Pavlovian-to-instrumental transfer, acetylcholine, tonically-active neurons, optogenetics, chemogenetics, biosensors, dopamine, incentive motivation

## Abstract

**Background:** Environmental reward-predictive stimuli provide a major source of motivation for adaptive reward pursuit behavior. This cue-motivated behavior is known to be mediated by the nucleus accumbens core (NAc). The cholinergic interneurons in the NAc are tonically active and densely arborized and, thus, well-suited to modulate NAc function. But their causal contribution to adaptive behavior remains unknown. Here we investigated the function of NAc cholinergic interneurons in cue-motivated behavior.

**Methods:** To do this, we used chemogenetics, optogenetics, pharmacology, and a translationally analogous Pavlovian-to-instrumental transfer behavioral task designed to assess the motivating influence of a reward-predictive cue over reward-seeking actions in male and female rats.

**Results:** The data show that NAc cholinergic interneuron activity is necessary and sufficient to oppose the motivating influence of appetitive cues. Chemogenetic inhibition of NAc cholinergic interneurons augmented cue-motivated behavior. Optical stimulation of acetylcholine release from NAc cholinergic interneurons prevented cues from invigorating reward-seeking behavior, an effect that was mediated by activation of β2-containing nicotinic acetylcholine receptors.

**Conclusions:** Thus, NAc cholinergic interneurons provide a critical regulatory influence over adaptive cue-motivated behavior and, therefore, are a potential therapeutic target for the maladaptive cue-motivated behavior that marks many psychiatric conditions, including addiction and depression.

---

Environmental reward-predictive stimuli provide a major source of motivation for adaptive reward pursuit behaviors (1). This incentive motivational value can become dysfunctional in many psychiatric disease states (2). Indeed, it can become amplified allowing cues to become potent triggers for maladaptive compulsive overeating (3), alcohol abuse (4–7), or drug seeking (8–12). Stress, anxiety, and depression (13–16) can also disrupt the motivating influence of appetitive cues, resulting in dampened or inappropriate motivation. The nucleus accumbens core (NAc) has been implicated in cue-motivated behavior (17–19). But little is known about the function of the major NAc neuromodulator acetylcholine. Such information is crucial given the purported importance of cholinergic signaling in many mental illnesses (20, 21).

Cholinergic interneurons provide the primary, though not exclusive (22), source of acetylcholine in the NAc (23). Despite comprising only 1-2% of the population, these large-bodied, tonically active neurons are densely arborized (24–29), making them ideally suited to modulate NAc function and associated behaviors. Cholinergic interneurons have also been shown to locally regulate striatal dopamine release (30–32). NAc cholinergic signaling is elevated under conditions that discourage vigorous reward seeking, such as satiety (33, 34), and has been implicated in anxiety- and depression-like states (35, 36) marked by blunted motivation. Cholinergic interneurons are transiently activated by cues signaling disadvantageous high effort, low reward conditions (37) or when movement must be withheld to obtain reward (38). Cues that encourage motivated behavior, such as those signaling reward availability, cause a characteristic pause in cholinergic interneuron activity (29, 37, 39–46). Yet still, very little is known of the causal contribution of NAc cholinergic interneurons to motivation.

We sought to fill this gap in knowledge by assessing the function of NAc cholinergic interneurons in cue-motivated behavior. Working from the evidence that cholinergic interneurons tend to increase their activity when vigorous motivated behavior is disadvantageous and pause when active reward pursuit is encouraged, we tested the hypothesis that NAc cholinergic interneuron activity functions to oppose the motivating influence of appetitive cues. Chemogenetic and optogenetic methods were used to selectively manipulate NAc cholinergic interneuron activity. We used the Pavlovian-to-instrumental transfer (PIT) test to measure cue-motivated behavior. This test is translationally analogous to that used in humans in health and disease (5, 11, 17, 47–55) and assesses the invigorating influence of an environmental reward-predictive stimulus over instrumental reward-seeking activity. Because the Pavlovian and instrumental components are independently trained, PIT isolates the incentive motivational value of the cue from other processes through which cues trigger action, such as via discriminative control or a stimulus-response relationship.

## MATERIALS AND METHODS

### Subjects

Adult (3-5 months) male and female ChAT::Cre+ transgenic rats (Long-Evans background) (56) were used for all experiments. Pups were weaned at PND 21 and group housed until experiment onset. Handling occurred daily, beginning at PND 60. Training and test were performed during the dark phase of a 12:12 hr reverse dark/light cycle. Rats were food-restricted to ~85% free-feeding body weight and water was provided *ad libitum* in the home cage. All procedures were conducted in accordance with the National Research Council’s Guide for the Care and Use of Laboratory Animals and were approved by the UCLA Institutional Animal Care and Use Committee.

### Surgery

Standard surgical procedures, described previously (57–59), were used for infusion of adeno-associated viruses (AAVs) and implantation of optical fiber or microinfusion injector/optical fiber guide cannula into the NAc core. Rats were anesthetized with isoflurane and a nonsteroidal anti-inflammatory agent was administered pre- and post-operatively to minimize pain and discomfort. Surgical details for each experiment are provided in the Supplemental Methods. Expression and placement was verified with standard histological procedures described in the Supplemental Methods.

### Behavioral Procedures

#### General training and testing

##### Training

Rats received Pavlovian and instrumental training in Med Associates conditioning chambers, as described previously (57, 60, 61).

##### Pavlovian conditioning

Rats first received 8 days of Pavlovian training in which 1 of 2 auditory stimuli (75 dB tone or white noise; counterbalanced across rats) was paired with non-contingent delivery of 45 mg chocolate-flavored, grain-based pellets (Bio-Serv, Frenchtown, NJ). During each 2-min presentation of the conditional stimulus (CS^+^), pellets were presented on a random time (RT)-30s schedule. The CS^+^ was presented 6x/session with a random 2-4 min inter-trial interval (mean=3 min). The lever was never present during these sessions.

##### Instrumental conditioning

All rats then received 8 days of instrumental training in which lever pressing earned delivery of a single chocolate pellet. Each session lasted until 20 outcomes had been earned, or 30 min elapsed. Rats received one day each of continuous, random interval (RI)-15s, and RI-30s schedules of reinforcement, followed by 5 days on the final RI-60s schedule. The CS^+^ was never present during this training.

##### CS^Ø^ habituation

Rats received 1 session of habituation to the neutral control stimulus (CS^Ø^), which consisted of 6, 2-min presentations of the CS^Ø^ (opposite stimulus as the CS^+^), with a 2-4 min inter-trial interval. No rewards were delivered during this session.

##### Pavlovian-to-instrumental transfer test

On the day prior to each PIT test, rats were given a single 30-min instrumental extinction session in which no cues were present, and the lever was available, but presses were unrewarded. During each PIT test the lever was continuously available, but pressing was not reinforced. Responding was extinguished for 5 min to establish a low rate of baseline performance, after which each CS was presented 4 times in pseudorandom order, also without accompanying reward. Each CS lasted 2 min with a 4-min fixed inter-trial interval. Rats received 1 Pavlovian and 2 instrumental retraining sessions identical to those above in between subsequent PIT tests. In all cases, testing commenced at least 4 weeks post-viral infusion to allow for construct expression.

#### Chemogenetic inactivation of NAc cholinergic interneurons

Prior to training, ChAT::Cre+ rats were bilaterally infused with a cre-inducible AAV vector to express the inhibitory designer receptor *human M4 muscarinic receptor* (hM4D(Gi)) or control fluorophore mCherry selectively in cholinergic interneurons of the NAc. Following training, rats received PIT tests, counterbalanced for order, one following vehicle and one following i.p. injection of the hM4D(Gi) ligand clozapine-N-oxide (CNO; 5mg/kg; see Supplemental Methods). These experiments were run in two separate cohorts and data were collapsed across cohorts following analyses indicating no interaction between Cohort and any of the variables of primary interest (hM4D(Gi): highest *F*=3.23, *P*=0.09; mCherry highest *F*=3.876, *P*=0.07). Final hM4D(Gi) *N*=19 (8 female; 2 rats were excluded due to off target viral spread) and mCherry *N*=16 (8 female). Following PIT testing, a subset of subjects were tested for the influence of NAc cholinergic interneuron inactivation on food consumption and lever pressing on a progressive ratio response requirement (see Supplemental Methods).

#### Optical stimulation of NAc cholinergic interneurons

Prior to training, ChAT::Cre+ rats were bilaterally infused with a cre-inducible AAV vector to express the excitatory opsin *channelrhodopsin-2* (ChR2) or control fluorophore eYFP selectively in NAc cholinergic interneurons of the NAc. Optical fibers were implanted bilaterally in the NAc. From the last 2 days of instrumental training and for a single additional Pavlovian retraining session, rats were tethered to the patchcord, but no light was delivered to allow habituation to the optical tether. Following training, rats received 4 PIT tests, counterbalanced for order, with intervening retraining. During each test, optical fibers were connected via ceramic sleeves to patch cords attached to a commutator. Blue light (473 nm, 10 Hz, 10 mW, 5 ms pulse width, 120 s duration; see also Supplemental Methods) was delivered for optical activation of ChR2-expressing NAc cholinergic interneurons. For the main experimental condition, light was delivered concurrent with each of the 4 CS^+^ presentations, with light and CS^+^ onset and offset synced. There were 3 separate control conditions: light delivered concurrent with each CS^Ø^ presentation, light delivered during the CS-free 2-min baseline period immediately prior to each CS^+^ presentation, or light delivered during the CS-free 2-min baseline period immediately prior to each CS^Ø^ presentation. There were no significant differences in performance between the preCS+ and preCS^Ø^ stimulation tests and, thus, data were collapsed across these tests into a single ‘baseline stimulation’ control condition (see Supplemental Figure 4). Final ChR2 *N*=9 (5 female; 5 subjects excluded for lack of expression and/or optical fiber misplacement), eYFP *N*=8 (5 female).

##### Optical stimulation of NAc cholinergic interneurons and inactivation of NAc β2-containing nAChRs

Prior to training, ChAT::Cre+ rats were bilaterally infused with a cre-inducible AAV vector to express ChR2 selectively in NAc cholinergic interneurons of the NAc. Microinfusion injector/optical fiber guide cannula were implanted bilaterally above the NAc. Following training, rats received 4 PIT tests, counterbalanced for order with intervening retraining. Prior to each test, rats were bilaterally infused with either the selective α4β2-containing nicotinic receptor competitive antagonist dihydro-β-erythroidine (DhβE; 15 μg/0.5 μl/side; see Supplemental Methods) or artificial cerebral spinal fluid (ACSF) vehicle via an injector inserted through the guide cannula designed to protrude 2.5 mm to just above the NAc (−6.5 mm). Following infusion, injectors were removed and optical fibers, also designed to protrude 2.5 mm and, thus target the NAc, were placed through guide cannula and secured via ceramic sleeves. During 2 of the tests, one each following vehicle or DhβE, blue light (473 nm, 10 Hz, 10 mW, 5 ms pulse width, 120 s duration) was delivered for optical activation of ChR2-expressing NAc cholinergic interneurons concurrent with each CS^+^ presentation. During the other two tests, an optical fiber was attached but no light was delivered. Thus, each rat received 4 tests: Vehicle/No stimulation, Vehicle/stimulation during CS^+^, DhβE/no stimulation, DhβE/stimulation during CS^+^. Following the PIT tests, optical fibers were removed and dummies were placed in the guide cannula. Final *N*=11 (all male, 1 rat was excluded due to a clogged cannula).

### Data analysis

#### Behavioral analysis

Lever pressing and entries into the food-delivery port were the primary behavioral output measures for the PIT test. These measures were counted for each 2-min CS period, with behavioral output during the 2-min periods prior to each CS serving as the baseline. For both the chemogenetic inhibition and optical stimulation experiments there was no interaction between trial and any of the other variables on lever pressing during the test (highest *F*=1.84, *P*=0.13). Thus, in all cases, data were collapsed across trials.

#### Sex differences

Approximately half the subjects in the chemogenetic and optical manipulation experiments were female. In neither case was there a main effect of Sex (hM4D(Gi): *F*_1,7_=2.72, *P*=0.12; ChR2: *F*_1,7_=0.71, *P*=0.43) and Sex did not significantly interact with the effect of CS and/or Drug or Stimulation period on lever pressing (highest *F*=3.41, *P*=0.08). Thus, all data were collapsed across sexes. Because sex did not influence results of the initial optogenetic experiment, the follow-up experiment assessing the influence of intra-NAc DhβE on the behavioral effect of optical stimulation included only males.

#### Statistical analysis

Data were processed with Microsoft Excel (Redmond, WA). Statistical analyses were conducted with GraphPad Prism (La Jolla, CA) and SPSS (IBM Corp, Chicago, IL). Data were analyzed with Student’s *t* tests, one-, two-, and three-way repeated-measures analysis of variance (ANOVA; Geisser-Greenhouse correction). Corrected post-hoc comparisons were used to clarify main effects and interactions. All datasets met equal covariance assumptions, justifying ANOVA interpretation (62). Alpha levels were set at *P*<0.05.

### Approach validation

Optical stimulation and chemogenetic inhibition of NAc cholinergic interneurons was validated *in vivo* with electroenzymatic choline biosensors and constant-potential amperometry as detailed in the Supplemental Methods. Briefly, to confirm chemogenetic inhibition of NAc cholinergic interneurons, silicon wafer-based platinum microelectrode array choline biosensors packaged with an optical fiber affixed to the back surface of the probe (to reduce the photovoltaic artifact) were lowered into the NAc of anesthetized rats expressing ChR2 and hM4D(Gi) in cholinergic interneurons. The ability of blue light (473 nm, 20 Hz, 5-30 mW, 10-ms pulse width, 5-s duration) to evoke acetylcholine release continuously monitored by the sensor was assessed following injection of vehicle or CNO (5 mg/kg, i.p.). Final *N*=4 recording locations in 2 subjects. To confirm stimulation of NAc cholinergic interneurons with the exact light parameters used in the behavioral experiments, choline biosensors/optical fibers were lowered into the NAc of anesthetized rats expressing ChR2 or eYFP in cholinergic interneurons. Choline fluctuations were monitored and blue light (473 nm, 10 Hz, 10 mW, 5-ms pulse width, 120-s duration) was delivered to evaluate its ability to evoke acetylcholine release in ChR2-expressing subjects. Final ChR2 *N*=5 recording locations in 4 subjects, eYFP *N*=5 recording locations in 3 subjects.

## RESULTS

### Chemogenetic inhibition of NAc cholinergic interneurons augments cue-motivated behavior

To evaluate the contribution of NAc cholinergic interneurons to cue-motivated behavior, we first chemogenetically inactivated these cells during a PIT test. Inactivation was achieved by using ChAT::Cre+ rats and a cre-inducible AAV vector to express the inhibitory designer receptor hM4D(Gi) selectively in cholinergic interneurons of the NAc (Figure 1A-C). CNO (5 mg/kg, i.p.) activation of hM4D(Gi) in cholinergic interneurons was found to effectively attenuate optically-evoked NAc acetylcholine release *in vivo* (Figure 1D).

**Figure 1.**
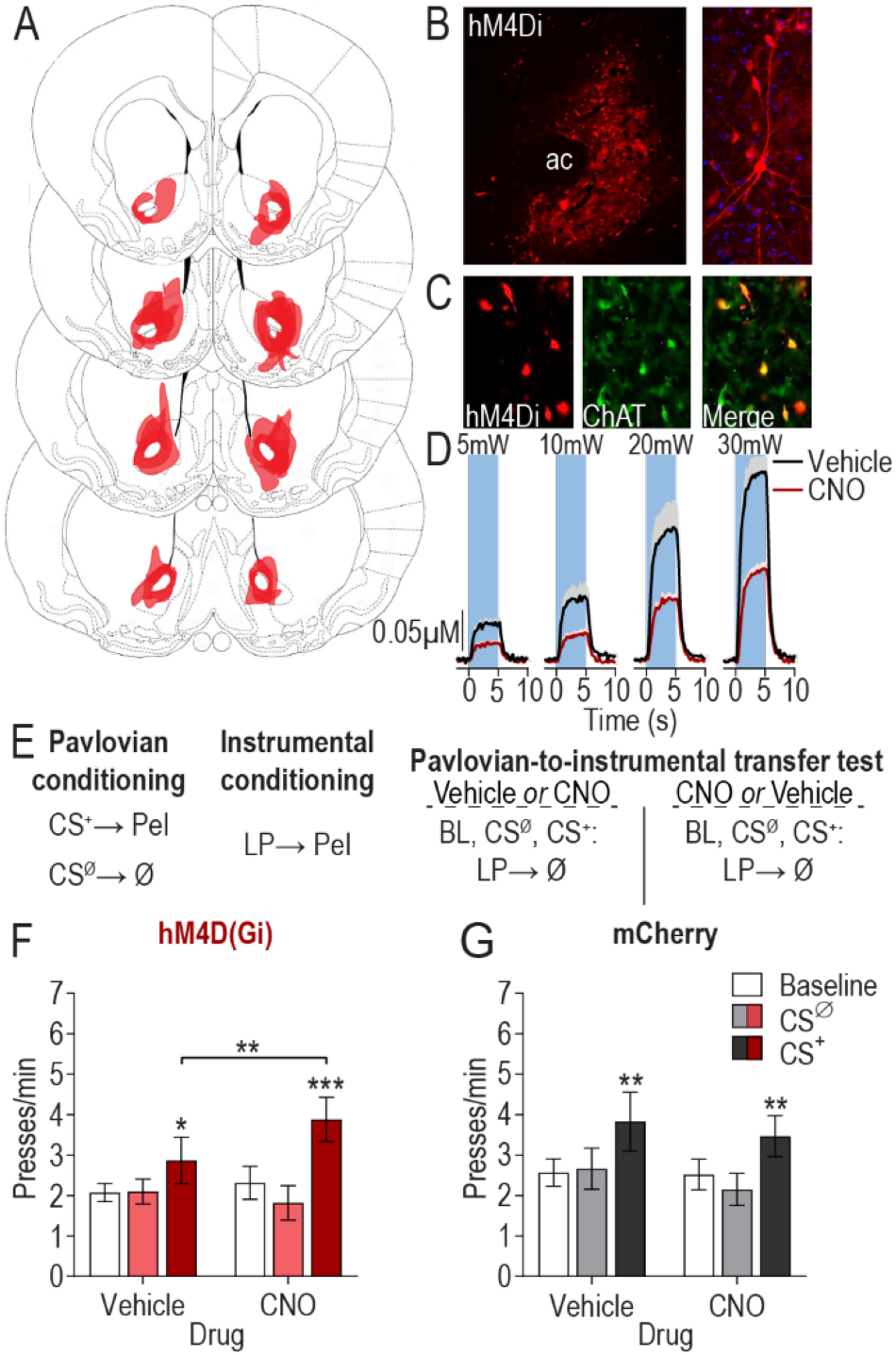
Chemogenetic inhibition of nucleus accumbens cholinergic interneurons augments cue-motivated behavior. (**A**) Schematic representation of hM4D(Gi)-mCherry expression in the NAc for all subjects. Slides represent 0.7 - 1.7 mm anterior to bregma. Images taken from (99). (**B**) Representative immunofluorescent images of hM4D(Gi)-mcherry expressing cholinergic interneurons in the NAc. AC, anterior commissure. (**C**) Colocalization of ChAT staining and hM4D(Gi)-mcherry expression in the NAc. (**D**) CNO:hM4D(Gi) attenuation of optically-evoked (473 nm, 20 Hz, 5-30 mW, 10-ms pulse width, 5-s duration) acetylcholine release in the NAc *in vivo* (see Supplemental Figure 1 for histology). Mean +1 s.e.m. (**E**) Procedure schematic. CS^+^, reward-predictive cue; CS^Ø^, neutral control stimulus; Pel, pellet reward; LP, lever press; Ø, no reward; Veh, Vehicle; CNO, Clozapine N-oxide. (**F-G**) Lever press rate during each 2-min period of the PIT test, averaged across trials compared between the CS-free (baseline), neutral stimulus (CS^Ø^), and reward-predictive cue (CS^+^) periods for the vehicle- and CNO-treated conditions in hM4D(Gi) (**F**) or mCherry control (**G**) subjects. Mean +1 s.e.m. \**P*<0.05, \*\**P*<0.01, \*\*\**P*<0.001.

Rats received Pavlovian training to pair a 2-min auditory conditional stimulus (CS^+^) with food pellet reward (Figure 1E). An alternate 2-min auditory stimulus was presented unpaired with reward and served as a control (CS^Ø^). Rats were then instrumentally conditioned, in the absence of the stimuli, to lever press to earn food rewards (see Supplemental Table 1 for training data). At the PIT test, the lever was available and each CS was presented in pseudorandom order to assess the motivating influence of the CS^+^ over lever-pressing activity. No rewards were delivered during this test. Increased lever-press rate during the CS^+^ provided the measure of cue-motivated behavior (i.e., expression of PIT). Each rat was tested twice, once following injection of vehicle and once following CNO, counterbalanced for order (Figure 1E).

Inactivation of NAc cholinergic interneurons augmented the expression of PIT (CS period: *F*_2,36_=8.15, *P*=0.001; Drug: *F*_1,18_=0.78, *P*=0.39; CS x Drug: *F*_2,36_=5.2, *P*=0.01; Figure 1F). Demonstrating PIT, the CS^+^ elevated lever pressing relative to both the baseline and CS^Ø^ periods under vehicle control conditions (*P*<0.05). Inactivation of NAc cholinergic interneurons enhanced the invigorating influence of the CS^+^ relative to the vehicle control condition (*P*<0.01). NAc cholinergic interneuron inactivation predominantly influenced CS^+^-invigorated responding; neither baseline, nor CS^Ø^ lever-press rate were significantly altered in the CNO condition (P>0.05). There was no effect of CNO on the expression of PIT in subjects lacking the hM4D(Gi) transgene (CS period: *F*_2,30_=4.47, *P*=0.02; Drug: *F*_1,15_=0.31, *P*=0.58; CS x Drug: *F*_2,30_=0.45, *P*=0.64; Figure 1G). Inactivation of NAc cholinergic interneurons did not alter the expression of Pavlovian conditional food-port approach responses during the PIT test. It also did not alter lever pressing during a progressive ratio test or basic food consumption (Supplemental Figure 2). Thus, inactivation of NAc cholinergic interneurons selectively enhanced the motivating influence of a reward-predictive cue over instrumental behavior.

### Optical stimulation of NAc cholinergic interneurons concurrent with reward cue presentation blunts cue-motivated behavior

The chemogenetic inactivation results suggest that NAc cholinergic interneurons function to oppose cue-motivated behavior. To further test this, we next evaluated the influence of stimulation of NAc cholinergic interneurons on expression of PIT. We used optical stimulation to provide temporal specificity. The excitatory opsin ChR2 was selectively expressed in NAc cholinergic interneurons (Figure 2A-C) of ChAT::Cre^+^ rats. Optical stimulation (473 nm, 10 Hz, 10 mW, 2 min) of these cells at a frequency in the upper range of their normal firing rate (63, 64) was found to increase acetylcholine release *in vivo*. This increase was restricted to the light-on period (*F*_2,8_=15.15, *P*=0.01) and did not occur in subjects lacking the ChR2 transgene (Figure 2D). Following Pavlovian and instrumental training, during the PIT test, we used a within-subject design to stimulate NAc cholinergic interneurons either concurrent with each 2-min CS^+^ presentation, or, in separate control tests, each CS^Ø^ presentation, or an equivalent number and duration of CS-free baseline periods (Figure 2E).

**Figure 2.**
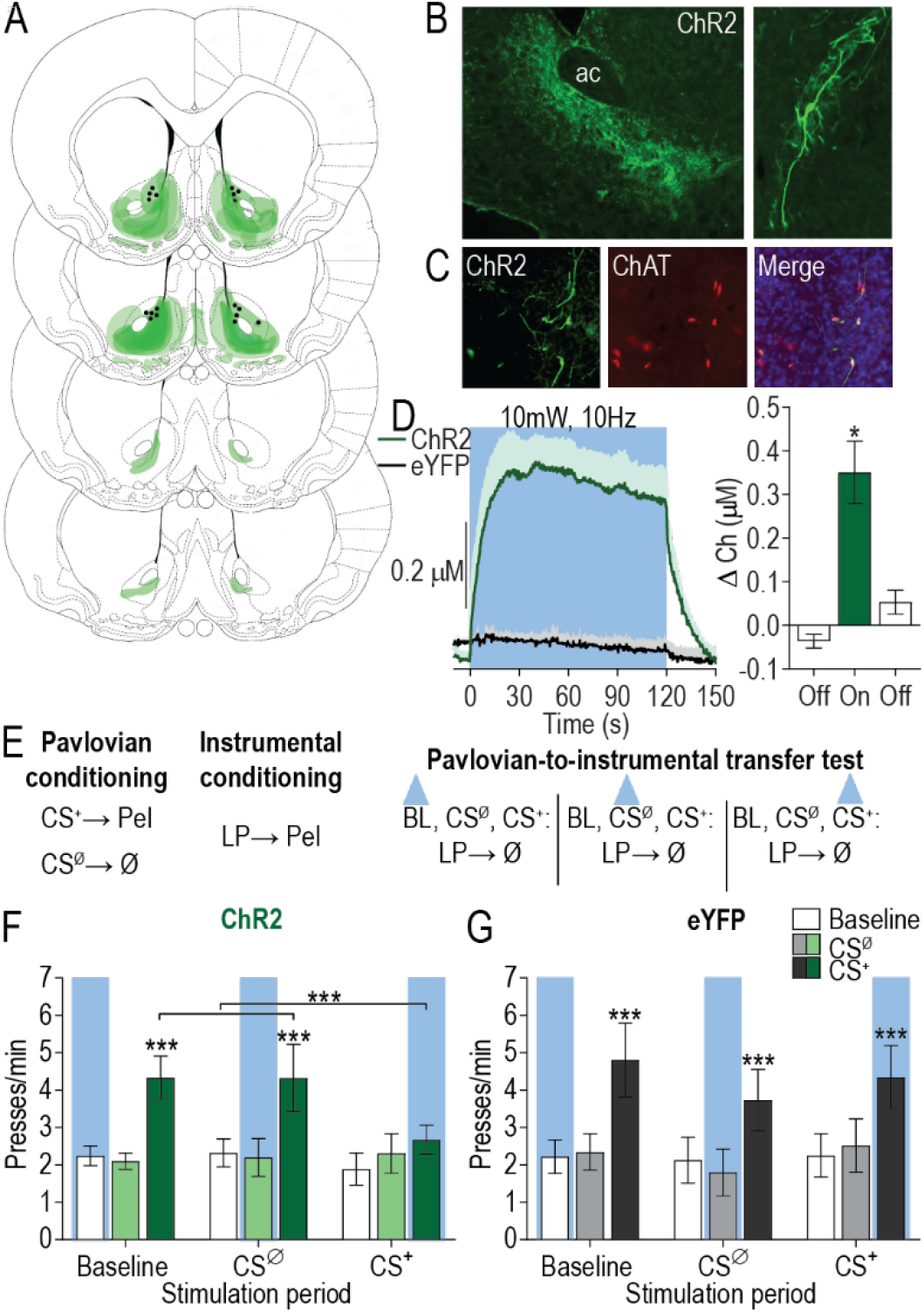
Optical stimulation of nucleus accumbens cholinergic interneurons concurrent with reward-predictive cue blunts cue-motivated behavior. (**A**) Schematic representation of ChR2-eYFP expression and fiber tips the NAc for all subjects. Slides represent 0.7 - 1.7 mm anterior to bregma. (**B**) Representative immunofluorescent images of ChR2-eYFP expressing cholinergic interneurons in the NAc. AC, anterior commissure. (C) Colocalization of ChAT staining and ChR2-eYFP expression in the NAc. (**D**) Optically-evoked acetylcholine release *in vivo* by blue light delivery (473 nm, 10 Hz, 10 mW, 5-ms pulse width, 120-s duration) to ChR2-expressing cholinergic interneurons in the NAc (see Supplemental Figure 3 for histology). Mean +1 s.e.m. (**E**) Procedure schematic. CS^+^, reward-predictive cue; CS^Ø^, neutral control stimulus; Pel, pellet reward; LP, lever press; Ø, no reward; blue triangle, light delivery. (**F-G**) Lever press rate during each 2-min period of the PIT test, averaged across trials compared between the CS-free (baseline), neutral stimulus (CS^Ø^), and reward-predictive cue (CS+) periods for tests in which optical stimulation occurred during the baseline stimulation, CS^Ø^, and CS^+^ periods in ChR2 (**F**) or eYFP control (**G**) subjects. Mean +1 s.e.m. \*\*\**P*<0.001.

Optical stimulation of NAc cholinergic interneurons during CS^+^ presentation blunted the expression of PIT (CS period: *F*_2,16_=8.07, *P*=0.004; Stimulation period: *F*_2,16_=0.71, *P*=0.50; CS x Stimulation period: *F*_4,32_=3.79, *P*=0.01; Figure 2F). Neither baseline nor CS^Ø^ period stimulation altered lever pressing during those periods (*P*>0.05) or the significant enhancement in such pressing induced by the CS^+^ (*P*<0.001). However, stimulation of NAc cholinergic interneurons concurrent with CS^+^ presentation prevented that cue from increasing lever pressing (*P*>0.05). Light delivery had no effect on the expression of PIT in subjects lacking the ChR2 transgene (CS period: *F*_2,14_=8.656, *P*=0.004; Stimulation period: *F*_2,14_=0.27, *P*=0.77; CS x Stimulation period: *F*_4,28_=1.04, *P*=0.41; Figure 2G). Optical stimulation of NAc cholinergic interneurons did not prevent the CS^+^ from eliciting Pavlovian conditional food-port approach responses (Supplemental Figure 5), suggesting no deficit in CS^+^ recognition. Thus, optical stimulation of NAc cholinergic interneurons blunted the expression of cue-motivated behavior.

### Acetylcholine release from NAc cholinergic interneurons works via β2-containing nicotinic receptors to blunt cue-motivated behavior

These data suggest that cholinergic interneuron activity tempers the motivating influence of reward-predictive cues over reward-seeking actions. Acetylcholine receptors are broadly distributed in the NAc and consist of two major subtypes: metabotropic muscarinic (mAChR) and ionotropic nicotinic (nAChR). We previously identified that activity of the NAc nAChRs, in particular, works to restrain the expression of cue-motivated behavior (57). Moreover, nAChRs containing the β2 subunit have been shown to be located exclusively on dopamine terminals (65) where they regulate phasic dopamine release (66–71), which has itself, in the NAc, been shown to track and mediate cue-motivated behavior (9, 60, 61, 72–75). Thus, we next asked whether the attenuating effect of optical stimulation of NAc cholinergic interneurons over cue-motivated behavior is mediated via these β2-containing nAChRs. To achieve this, we again selectively expressed ChR2 in NAc cholinergic interneurons (Figure 3A-C) and evaluated the influence of intra-NAc infusion of dihydro-β-erythroidine (DhβE; 15μg/side), a selective α4β2-containing nAChR antagonist, on the suppressive influence of NAc cholinergic interneuron stimulation over PIT expression (Figure 3D).

**Figure 3.**
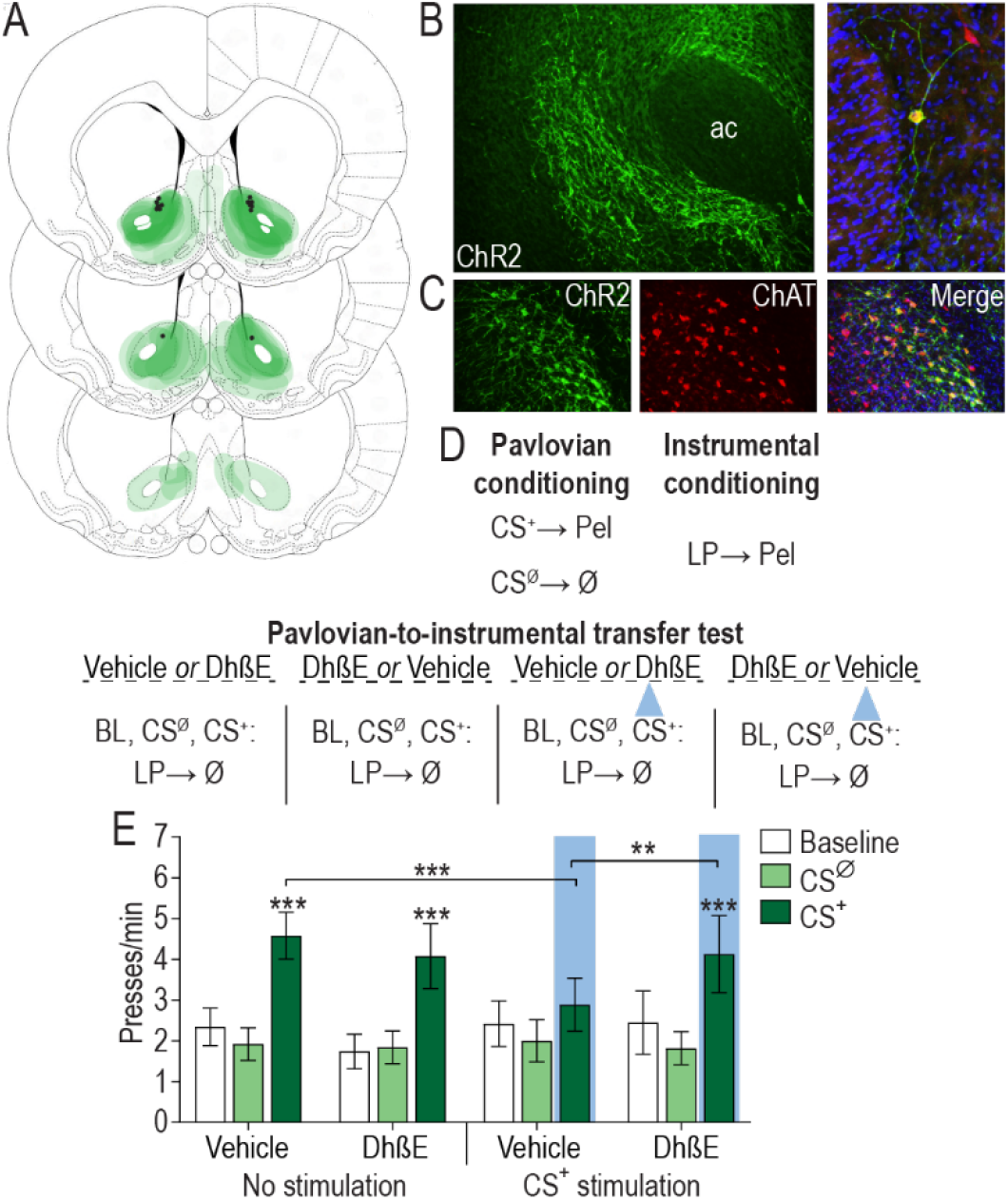
Acetylcholine release from nucleus accumbens cholinergic interneurons works via β2-containing nicotinic receptors to blunt cue-motivated behavior. (**A**) Schematic representation of ChR2-eYFP expression and fiber/injector tips the NAc for all subjects. Slides represent 0.7 - 1.7 mm anterior to bregma. (**B**) Representative immunofluorescent images of ChR2-eYFP expressing cholinergic interneurons in the NAc. AC, anterior commissure. (**C**) Colocalization of ChAT staining and ChR2-eYFP expression in the NAc. (**D**) Procedure schematic. CS^+^, reward-predictive cue; CS^Ø^, neutral control stimulus; Pel, pellet reward; LP, lever press; Ø, no reward; Veh, Vehicle; DhβE, Dihydro-β-erythroidine; blue triangle, light delivery. (**E**) Lever press rate during each 2-min period of the PIT test, averaged across trials compared between the CS-free (baseline), neutral stimulus (CS^Ø^), and reward-predictive cue (CS+) periods for the tests with either intra-NAc vehicle or DhβE with or without optical stimulation during the CS^+^. Mean +1 s.e.m. \*\**P*<0.01, \*\*\**P*<0.001.

Blockade of β2-containing nAChRs recovered the impairment of PIT induced by optical stimulation of NAc cholinergic interneurons during the CS^+^ (CS period: *F*_2,22_=22.69, *P*<0.0001; Optical stimulation: *F*_1,11_=0.082, *P*=0.78; Drug: *F*_1,11_=0.003, *P*=0.96; CS x Stimulation: *F*_2,22_=5.19, *p*=0.02; CS x Drug x Stimulation: *F*_2,22_=5.10, *P*=0.02; Figure 3E). We replicated the suppressive effect of optical stimulation of NAc cholinergic interneurons during CS^+^ presentation on the expression of PIT relative to a non-stimulated control condition (*P*>0.001). Whereas intra-NAc infusion of DhβE alone at this dose did not influence PIT expression relative to the vehicle-infused control condition (*P*>0.05), it did alleviate the suppressive effect of cholinergic interneuron stimulation (*P*<0.01), allowing subjects to show a significant PIT effect (*P*<0.001). These data demonstrate that acetylcholine release from NAc cholinergic interneurons acts via β2-containing nAChRs to blunt the motivating influence of cues. Secondarily, they indicate that the effect of optical stimulation of cholinergic interneurons was not due to nAChR receptor desensitization.

## DISCUSSION

Using a combination of chemogenetic, optogenetic, and pharmacological approaches, we investigated the function of NAc cholinergic interneurons in cue-motivated behavior. The data revealed that cholinergic interneuron activity in the NAc functions to limit the motivational influence of reward-predictive cues over reward-seeking actions. Chemogenetic inactivation of NAc cholinergic interneurons augmented cue-motivated behavior, whereas optical stimulation of these cells temporally restricted to cue presentation prevented cues from motivating action. This mitigating function is achieved via acetylcholine activation of β2-containing nAChRs.

These data accord well with evidence of the activity patterns of striatal cholinergic interneurons collected in non-human primates and rodents. Striatal cholinergic interneurons can both tonically and phasically increase their activity when vigorous motivated behavior is discouraged, for example in states of satiety (33, 34), or when cues signal unfavorable (e.g., high effort, low reward) conditions (37). Cholinergic interneurons also transiently increase their activity when cues signal that reward is available contingent upon a no-go response (38), i.e., when motivated movement must be withheld. Striatal cholinergic interneurons transiently pause their activity in response to cues signaling that vigorous reward seeking is advantageous. For example, cholinergic interneurons will pause in response to reward-predictive cues (29, 37, 39–46) and when cues signal favorable low effort/high reward conditions (37). The current data provide an important causal addition to this literature and reveal that increases in NAc cholinergic interneuron activity function to oppose cue-motivated behavior and that decreases or pauses in such activity are permissive to cue-motivated action. These results also indicate that the NAc inputs that regulate cholinergic interneuron excitability, activity, or synchrony, such as thalamostriatal projections (68), are well-positioned to influence cue-motivated behavior.

We found the suppressive effect of optical stimulation over cue-motivated behavior to depend on activity of β2-containing nAChRs. Acetylcholine release from NAc cholinergic interneurons acts at β2-containing nAChRs receptors to curtail the motivating influence of appetitive cues. This is consistent with our previous evidence that general nAChR, but not mAChR, blockade augments cue-motivated behavior (57). Moreover, that inactivation of β2-containing nAChRs completely recovered the suppressive influence of optical stimulation of NAc CINs over cue-motivated behavior, suggests that, although other acetylcholine receptor subtypes may contribute, β2-containing nAChR receptors are a critical locus of action for cholinergic regulation of cue-motivated behavior.

NAc core dopamine release is a major substrate of cue-motivated behavior. Its activity correlates with (57, 60, 73, 76) and is necessary (61, 75, 77) and sufficient (74, 78, 79) for the motivational influence of reward-predictive cues. β2-containing nAChRs are located exclusively on DA terminals (65), where they have been found to terminally modulate dopamine release (66–71). The present data may be considered surprising in light of evidence that optical stimulation of striatal cholinergic interneurons can evoke dopamine release from terminals via action at β2-containing nAChRs (68, 69). But a growing body of literature indicates that cholinergic regulation of dopamine release depends on the activity state of the dopamine cells (80, 81). β2-containing nAChR activity facilitates low probability (32, 66, 82) and tonic dopamine release (83), but will actually suppress dopamine release that results from high-frequency stimulation, which mimics dopamine neuron burst firing (32, 66, 82). Indeed, inactivation of β2-containing nAChRs in the NAc will augment dopamine release induced by high frequency stimulation *ex vivo* (67, 84) and general nAChR inactivation in the NAc will potentiate the phasic dopamine release response to reward-predictive cues in awake-behaving animals (57). Thus, we speculate that NAc cholinergic interneuron activity may restrain the motivating influence of reward-predictive cues via attenuating their ability to elicit dopamine release, with pausing in their signaling being permissive to such release and associated motivation.

The suppressive function of NAc cholinergic interneurons over cue-motivated behavior is interesting in light of how these cells are regulated. NAc cholinergic interneurons are controlled by several factors that mediate food-related motivation and responsivity to food cues. For example, they express receptors for the adiposity and satiety signal insulin, activation of which increases their activity and modulates NAc dopamine signaling through a nAChR-dependent mechanism (85). They also express receptors for corticotropin releasing factor (CRF), which mediates the positive and negative effects of stress (86–89). NAc CRF receptor activation increases cholinergic interneuron activity (90) and acetylcholine release (91), and regulates dopamine release (90). Moreover, serotonin, a neuromodulator long linked to motivation and mood, and recently in the NAc linked to adaptive social behavior (92), attenuates the excitability of NAc cholinergic interneurons via presynaptic 5-HT1A and postsynaptic 5-HT1B receptors (93).

Thus, NAc cholinergic interneurons are well-positioned to mitigate cue-motivated behavior when vigorous motivated action would not be beneficial and to promote cue-motivated behavior when it is adaptive. Dysfunction in this mechanism could, therefore, lead to the dysregulated motivation underlying some mental illnesses. Indeed, cues can become unnaturally strong motivators of drug-seeking behavior in addiction (4, 8, 94, 95) and NAc cholinergic interneurons have been linked to addiction-like behaviors (96, 97). Depression can be characterized by avolitional symptoms (94, 98), and NAc cholinergic interneurons have been linked to depression-like behavior (35). These results, therefore, have implications for the understanding and treatment of these and other diseases marked by maladaptive motivation.

## ACKNOWLEDGEMENTS

This research was supported by NIH grants MH106972 to KMW and SBO, AG045380, DK098709, and DA029035 to SBO, DA035443 to KMW, NS087494 to HGM and KMW, and T32 DA024635 to ALC. It was also supported by a UCLA dissertation year fellowship to ALC and a UCLA Integrated and Interdisciplinary Undergraduate Research Program Fellowship to TJA. The authors would like to thank Dr. Melissa Malvaez for helpful comments on the manuscript and Ana Sias for her assistance with histology.

## AUTHOR CONTRIBUTIONS

ALC, KMW, and SBO conceptualized and designed the experiments and interpreted the data. ALC and KMW analyzed the data. ALC conducted the optogenetic experiments and optogenetic validation, with assistance from VYG. TJA and ALC conducted the chemogenetic experiments, with assistance from VYG. CS and KMW conducted the chemogenetic validation experiments. IH, HGM, and KMW designed the choline biosensors and IH prepared and tested all sensors. KMW and ALC wrote the manuscript with assistance from TJA and SBO.

## SUPPLEMENTAL MATERIAL

### SUPPLEMENTAL MATERIALS AND METHODS

#### Surgery

Standard surgical procedures described previously (57) were used for all surgeries. Rats were anesthetized with isoflurane (4–5% induction, 1–2% maintenance) and a nonsteroidal anti-inflammatory agent was administered pre- and post-operatively to minimize pain and discomfort. Following surgery rats were individually housed. Virus was infused into the NAc core at a rate of 0.1μl/min via an infusion needle. Following infusion, the injectors were left in place for 10 min, then slowly removed to prevent undesired viral spread.

##### Chemogenetic inhibition of NAc cholinergic interneurons

Rats were randomly assigned to a viral group prior to onset of behavioral training and were infused bilaterally with adeno-associated virus (AAV) cre-dependently encoding the inhibitory designer receptor *human M4 muscarinic receptor* (hM4D(Gi)); AAV2-hSyn-DIO-hM4D(Gi)-mcherry; Titer 3.7 x 10^12^; Cohort 1: UNC-CH Vector Core, Chapel Hill, NC; Cohort 2: Addgene, Cambridge, MA) or control fluorophore only (AAV2-hSyn-DIO-mcherry; titer 5.6 x 10^12^ particles/ml; UNC-CH Vector Core). Virus (0.8μl) was infused into the NAc core (AP: +1.5 mm, ML: +/− 3.0, DV: −6.5, at a 9 degree angle).

##### Optical stimulation of NAc cholinergic interneurons

Rats were randomly assigned to a viral group prior to onset of behavioral training and were infused bilaterally with AAV cre-dependently expressing the excitatory opsin *channelrhodopsin-2* (ChR2; AAV5-EF1a-DIO-ChR2-eYFP; titer 4.8-7 x 10^12^ particles/ml; UNC-CH Vector Core) or control fluorophore only (AAV5-EF1a-DIO-eYFP; titer 6.5 x 10^12^ particles/ml, UNC-CH Vector Core). Virus was infused at 3 separate coordinates (AP: +1.3 mm, ML: +1.3, V: −6.8; AP: +1.3 mm, ML: +1.3, V: −6.0; AP: +1.3 mm, ML: +3.0, V: −6.5, at a 9° angle; 1 μl/infusion site; (57, 100)). At the same surgery, custom-, in-house-made optical fibers were implanted targeted at the NAc core (AP: +1.3 mm; ML: +1.3; V: −6.5). Prior to surgery, all optical fibers were tested for loss of power and only optical fibers with a loss of power >5% were used.

##### Optical stimulation of NAc cholinergic interneurons with inactivation of NAc β2-containing nAChRs

Prior to behavioral training rats were infused bilaterally with AAV cre-dependently expressing ChR2 as above. Custom-made microinfusion injector/optical fiber guide cannula (Doric Lenses, Quebec, QC, Canada) were implanted bilaterally targeted above the NAc core (AP: +1.3 mm; ML: +1.3; V: −4.0).

##### Validation of chemogenetic manipulation of NAc cholinergic interneurons

Rats received 2 separate surgeries, spaced 5 days apart. In one, rats were infused unilaterally with AAV cre-dependently expressing ChR2. In the other, rats were unilaterally in the same hemisphere with AAV cre-dependently expressing hM4D(Gi). One rat received the ChR2 AAV first and hM4D(Gi) second, and the other received the hM4D(Gi) AAV first. In each case, AAV was infused at 3 separate coordinates (AP: +1.3 mm, ML: +1.3, V: −6.8; AP: +1.3 mm, ML: +1.3, V: −6.0; AP: +1.3 mm, ML: +3.0, V: −6.5, at a 9° angle; 1 μl/infusion site; (57, 100)).

##### Validation of optical stimulation of nucleus accumbens cholinergic interneurons

Rats were infused bilaterally with AAV cre-dependently expressing ChR2 or the control fluorophore eYFP at 3 separate coordinates (AP: +1.3 mm, ML: +1.3, V: −6.8; AP: +1.3 mm, ML: +1.3, V: −6.0; AP: +1.3 mm, ML: +3.0, V: −6.5, at a 9° angle; 1 μl/infusion site; (57, 100)).

#### Behavioral training and testing

##### Apparatus

Training and testing took place in Med Associates conditioning chambers (East Fairfield, VT) housed within sound- and light-attenuating boxes, described previously (101). Behavioral testing for chemogenetic manipulations also occurred in these chambers. For optogenetic experiments, behavioral tests were conducted in Med associates conditioning chambers outfitted with an Intensity Division Fiberoptic Rotary Joint (Doric Lenses) connecting the output fiberoptic patchcords to a laser (Dragon Lasers, ChangChun, JiLin, China) positioned outside the conditioning chamber.

All chambers contained a retractable lever that could be inserted to the left of a recessed food-delivery port in the front wall. A photobeam entry detector was positioned at the entry to the food port. The chambers were equipped with a pellet dispenser that delivered a single 45-mg chocolate pellet (Bio-Serv, Frenchtown, NJ) into the food port. Both a tone (1.5 kHz) and white noise generator were attached to individual speakers on the wall opposite the lever and food-delivery port. A 3-watt, 24-volt house light mounted on the top of the back wall opposite the food-delivery port provided illumination.

##### Chemogenetic inactivation of NAc cholinergic interneurons prior to consumption test

Following PIT testing (see main text), hM4D(Gi)-expressing subjects were tested for the influence of NAc cholinergic interneuron inactivation on food consumption. All subjects were habituated to the feeding chambers (home cages devoid of bedding). After this, they were given 2, 1-hr test sessions in which they had free access to chocolate pellets (the food paired used for conditioning) following an injection of vehicle or CNO, counterbalanced for order. A subset of hM4D(Gi)-expressing subjects (*N*=8) also received a set of tests for consumption of home chow. A subset (*N*=9) of the mCherry-control subjects also received these consumption tests. Food was weighed prior to an after the consumption test to quantify amount consumed.

##### Chemogenetic inactivation of NAc cholinergic interneurons prior to progressive ratio test

Following PIT testing, a subset of subjects (hM4D(Gi) *N*=8; mCherry *N*=9) were shifted to a progressive ratio schedule of reinforcement for which lever pressing earned a chocolate pellet and after each earned reinforcer the response requirement increased by 50% (rounded up, beginning with a single press requirement). The session ended when 3 min elapsed without a lever press. Breakpoint was recorded as the highest ratio achieved. Rats received 3 days of progressive ratio training prior to 2 progressive ratio tests, with intervening retraining, one each following vehicle or CNO, order counterbalanced.

#### Drugs

##### Clozapine-N-oxide

For the first cohort of hM4D(Gi) and mCherry subjects, Clozapine-N-oxide (CNO) was obtained from Tocris (Minneapolis, MN) and was dissolved in Dimethyl-sulfoxide (DMSO; Sigma-Aldrich, St. Louis, MO), then diluted with 0.9% saline for a final 5% concentration of DMSO and a 5mg/mL concentration of CNO. Vehicle consisted of 5% DMSO in a 0.9% saline solution. For the second cohort, water soluble CNO was obtained from Enzo Life Sciences (Farmingdale, NY) and dissolved in sterile saline vehicle. In both cases, CNO was given at a 5mg/kg (102). Vehicle and CNO were injected i.p. at 1 ml/kg 30 minutes prior to test.

##### Dihydro-β-erythroidine

The selective α4β2-containing nicotinic receptor antagonist Dihydro-β-erythroidine (DhβE) was obtained from R&D systems (Minneapolis, MN), dissolved in sterile ACSF and infused as described previously (57, 103) bilaterally into the NAc at 15 μg/0.5 μl/side over 1 min. Given our previous work demonstrating that bilateral general blockade of nicotinic acetylcholine receptors with mecamylamine can augment cue-motivated behavior (57), we selected a subthreshold dose of DhβE that would not on its own influence motivated behavior when infused into the NAc (104). This avoided the confound of an independent effect of the drug when it was paired with optical stimulation of NAc cholinergic interneurons.

#### Optical stimulation

Light was delivered to the NAc using a laser (Dragon Lasers, ChangChun, JiLin, China) connected through a ceramic mating sleeve (Thorlabs, Newton, NJ) to the ferrule implanted on the rat. We used a 473 nm laser to activate ChR2-transfected projection neurons. For optical stimulation of NAc cholinergic interneurons in behaving subjects, blue (473 nm) light pulses (5 msec pulse width, 10 mW) were delivered at 10 Hz for 120s duration. Stimulation parameters were selected to match the upper range of the endogenous firing rate of striatal cholinergic interneurons (63, 64) and were found to reliably elicit acetylcholine release in the NAc *in vivo* (Figure 2D). Light effects were estimated to be restricted to the NAc core based on predicted irradiance values (https://web.stanford.edu/group/dlab/cgi-bin/graph/chart.php).

We also assess the influence of chemogenetic inhibition of NAc cholinergic interneurons over optically-evoked acetylcholine release. In this case, blue light pulses blue (473 nm) light pulses of varying magnitudes (10 msec pulse width, 5-30 mW) were delivered at 20 Hz for 5-s duration to evoke transient acetylcholine release.

#### Choline biosensors

##### Biosensor fabrication

Silicon wafer-based platinum microelectrode array (MEA) probes were fabricated in the Nanoelectronics Research Facility at UCLA as described previously (101, 105, 106) and modified for choline detection using a method similar that we have described for glutamate detection (59, 101, 105, 106). These biosensors use choline oxidase (ChOx) as the biological recognition element for choline and rely on electro-oxidation, via constant-potential amperometry (0.7 V versus a Ag/AgCl reference electrode), of enzymatically-generated hydrogen peroxide reporter molecule to provide a current signal. This current output is recorded and converted to choline concentration using a calibration factor determined *in vitro* prior to sensor implantation. Choline sensing allows for an accurate proxy measure of extracellular acetylcholine, which is rapidly hydrolyzed by endogenous acetylcholinesterase (107–110). Indeed, adding acetylcholinesterase onto the sensing electrode does not enhance detection of cholinergic activity (108). Enzyme immobilization was accomplished by manually loading enzyme mixture consisting of ChOx (4 μl of 0.5 u/μl ChOx) and bovine serum albumin (2 μl of 60mg/ml BSA) on the microelectrode sites, following by topping a layer of bis(sulfosuccinimidyl)suberate (100mg/ml BS3) for crosslinking. Interference from electroactive anions and cations is effectively excluded from the amperometric recordings, while still maintaining a subsecond response time, by electropolymerization of m-phenylenediamine (m-PD), as well as dip-coat application of Nafion to the electrode sites prior to enzyme immobilization. Electrodes were coated with 5 mM m-PD at 850 mV, followed by dip-coating in 2% Nafion, and cured at 115° C for 20 min. Each MEA had two non-enzyme-coated sentinel electrodes for the removal of correlated noise from the choline sensing electrodes by signal subtraction, as described previously (101, 106). These electrodes were prepared identically with the exception that the BSA solution did not contain ChOx.

##### Reagents

Choline oxidase (ChOx, from Alcaligenes sp), Nafion (5 wt.% solution in lower aliphatic alcohols/H2O mix), bovine serum albumin (BSA, min 96%), m-phenylenediammine (m-PD), choline chloride (>99%), L-ascorbic acid, 3-hydroxytyramine (dopamine) were purchased from Sigma (Sigma-Aldrich Co., St. Louis, MO, USA). Bis(sulfosuccinimidyl)suberate (BS3) was purchased from Thermo Fisher Scientific (Pittsburgh, PA).

##### Instrumentation

Electrochemical preparation of the sensors was performed using a Versatile Multichannel Potentiostat (model VMP3) equipped with the ‘p’ low current option and low current N’ stat box (Bio-Logic USA, LLC, Knoxville, TN). *In vitro* calibration and *in vivo* measurements were conducted using a low-noise multichannel Fast-16 mkIII potentiostat (Quanteon LLC, Nicholasville, KY), with reference electrodes consisting of a glass-enclosed Ag/AgCl wire in 3 M NaCl solution (for *in vitro*, Bioanalytical Systems, Inc., West Lafayette, IN) or a 200 μm diameter Ag/AgCl wire (*in vivo*). All potentials are reported versus the Ag/AgCl reference electrode. Oxidative current was recorded at 80 kHz and averaged over 0.25-s intervals.

##### In Vitro Biosensor Characterization

All biosensors were calibrated *in vitro* to test for sensitivity and selectivity to choline prior to implantation. A constant potential of 0.7 V was applied to this working electrodes against a Ag/AgCl reference electrode in 40 ml of stirred PBS at pH 7.4 and 37° C within a faraday cage. After the current detected at the electrodes equilibrated (~15-30 min), aliquots of choline were added to the beaker to reach final concentrations in the range of 20-100 μM choline. A calibration factor based on analysis of these data was calculated for each electrode. The average calibration factor the sensors used in these studies was 37.42 (s.e.m.=6.77) μM/nA. Control electrodes, coated with P(m-PD), Nafion, and BSA/BS3, but not ChOx, showed no detectable response to choline. Aliquots of ascorbic acid (250 μM final concentration) and dopamine (5-10 μM final concentration) were added to the beaker as representative examples of readily oxidizable potential anionic and cationic interferent neurochemicals, respectively, to confirm selectivity for choline. For the sensors used in these studies, no current changes above the level of the noise were detected to the addition of cationic (dopamine) or anionic (ascorbic acid) interferents, as reported previously (101, 111). To assess uniformity of H2O2 sensitivity across control and ChOx-coated electrodes, aliquots of H2O2 (10 μM) were also added to the beaker. There was less than a 10% difference in the H2O2 sensitivity on control electrode sites relative to enzyme-coated sites, indicating that any changes detected *in vivo* on the enzyme-coated biosensor sites following control channel signal subtraction could not be attributed to endogenous H2O2.

#### Approach validation

##### Validation of chemogenetic inhibition of cholinergic interneurons

Pre-calibrated, silicon wafer-based platinum microelectrode array choline biosensors were packaged with an optical fiber affixed to the back surface of the sensor (to reduce the photovoltaic artifact) such that the tip of the optical fiber was ~100 μm above the sensor tip. Rats expressing ChR2 and hM4D(Gi) in cholinergic interneurons of the NAc were anesthetized with isoflurane (induced at 5% and maintained at 1-2%) and placed in a stereotaxic frame. Breaths per minute (bpm) were maintained at 1 bpm by adjusting isoflurane level. The biosensor/optical fiber probe was lowered into the NAc core (AP: +1.3 mm; ML: +1.3; V: −6.5 - 6.8) and connected to the potentiostat for application of a 0.7 V potential. Oxidative current was recorded at 80 kHz and averaged over 0.25 s intervals.

Following equilibration of the amperometric signal (~30 min) laser stimulations were delivered via the optical fiber to evoke choline fluctuations and sensor placement was optimized along the DV axis to a recording location with reliable evoked choline fluctuations. Rats were then injected with sterile saline vehicle. Blue light (473 nm, 20 Hz, 5-30 mW, 10-ms pulse width, 5-s duration) was delivered to evoke acetylcholine release. Stimulations were spaced at least 30 s apart and the average current response across a 1-s period, 5 s prior to stimulation onset served as the baseline. Following completion of the stimulation protocol, rats were given CNO i.p., as described for the behavioral experiments above. 45 min later the stimulation protocol was repeated to determine the extent to which CNO activation of hM4D(Gi) attenuated optically-evoked acetylcholine release. Final ChR2+hM4D(Gi) *N*=4 recording locations in 2 subjects.

##### Validation of optical stimulation of cholinergic interneurons

Rats expressing ChR2 or eYFP in cholinergic interneurons of the NAc were anesthetized and implanted with a biosensor/optical fiber probe into the NAc (as above). Amperometric signal was recorded and sensor placement optimized as above. Blue light (473 nm, 10 Hz, 10 mW, 5-ms pulse width, 120-s duration) was delivered at the exact parameters used for the behavioral experiment to evaluate its ability to evoke acetylcholine release in ChR2-expressing subjects. Stimulations were spaced at least 60 s apart and the average current response across a 1-s period, 10 s prior to stimulation onset served as the baseline. Final ChR2 *N*=5 recording locations in 4 subjects, eYFP *N*=5 recording locations in 3 subjects.

#### Histology

Rats were deeply anesthetized with Pentasol (390mg/mL; 100mg/kg) and transcardially perfused with PFA. Brains were removed and post-fixed in 4% formalin for 20 hours, after which they were placed in 30% sucrose solution for 2-3 days. Brains were then frozen and sliced on a cryostat into 40-μm sections and stored in PBS.

We used ChAT immunoreactivity to visualize cholinergic interneurons. Sections were washed in PBS and blocked in 3% normal donkey serum with 0.3% triton X-100 for 2 hours at room temperature. Following washes in PBS, the tissue was incubated in goat-anti-ChAT (1:200; Millipore Sigma, Burlington, MA) at 4° Celsius for 20 hrs. It was then washed in PBS and incubated for 2 hrs at room temperature with either donkey-anti-goat RHD red (1:400; Millipore Sigma) or donkey anti-goat Cy2 (1:400; Abcam, Cambridge, UK).

To verify ChR2-eYFP expression, tissue sections were washed in PBS and blocked in 3% normal goat serum and 0.5% triton X-100 for 1.5 hrs. Following 3 PBS washes, tissue was incubated with mouse anti-GFP (1:1000, Abcam) 3% NGS and 0.5% triton X-100 at 4° Celsius for 20 hrs. Following 3 PBS washes, sections were incubated in goat anti-mouse Alexa 488 (1:1000, Abcam).

To verify hM4D(Gi)-mCherry expression, tissue sections were washed in PBS and blocked in 10% normal goat serum and 0.5% triton X-100 for 1.5 hrs. Following 3 washes at 5 min each, tissue was incubated with rabbit anti-mCherry (1:1000, ThermoFisher Scientific, Waltham, MA) at 4° C for 20 hrs. Following 3 blocking washes at 10 min each, tissue was incubated for 2 hrs with goat anti-rabbit alexa fluor 594 (1:500, ThermoFisher Scientific).

All sections were mounted with ProLong Gold antifade reagent with DAPI (Invitrogen, Carlsbad, CA) and imaged using a Keyence (BZ-X710) microscope (Osaka, Osaka Prefecture, Japan) with a 4X, 10X, or 20X objective (CFI Plan Apo), CCD camera, and BZ-X Analyze software. Tiled images were taken of whole sections, including 6-8 sections per rat containing the NAc across the A/P axis based on the Paxinos and Watson brain atlas (99).

#### Data analysis

##### Progressive ratio behavioral analysis

For the progressive ratio tests, lever press rate (presses/min) was calculated for the entire session and breakpoint was calculated as the highest ratio achieved before a >3-min break in lever pressing.

##### Amperometric analysis

Analysis details and characterization of release events have been described previously (59, 101, 106, 112). Electrochemical data were baseline-subtracted. Detected current was averaged across a 1-s period 5 or 10 s (as above) prior to stimulation and was subtracted from current output at each time point. Current changes from baseline on the PPD/Nafion-coated sentinel electrode were then subtracted from current changes on the PPD/Nafion/ChOx choline biosensor electrode to remove correlated noise. This signal was then converted to choline concentration using an electrode-specific calibration factor obtained *in vitro*.

**Supplemental Table 1.**
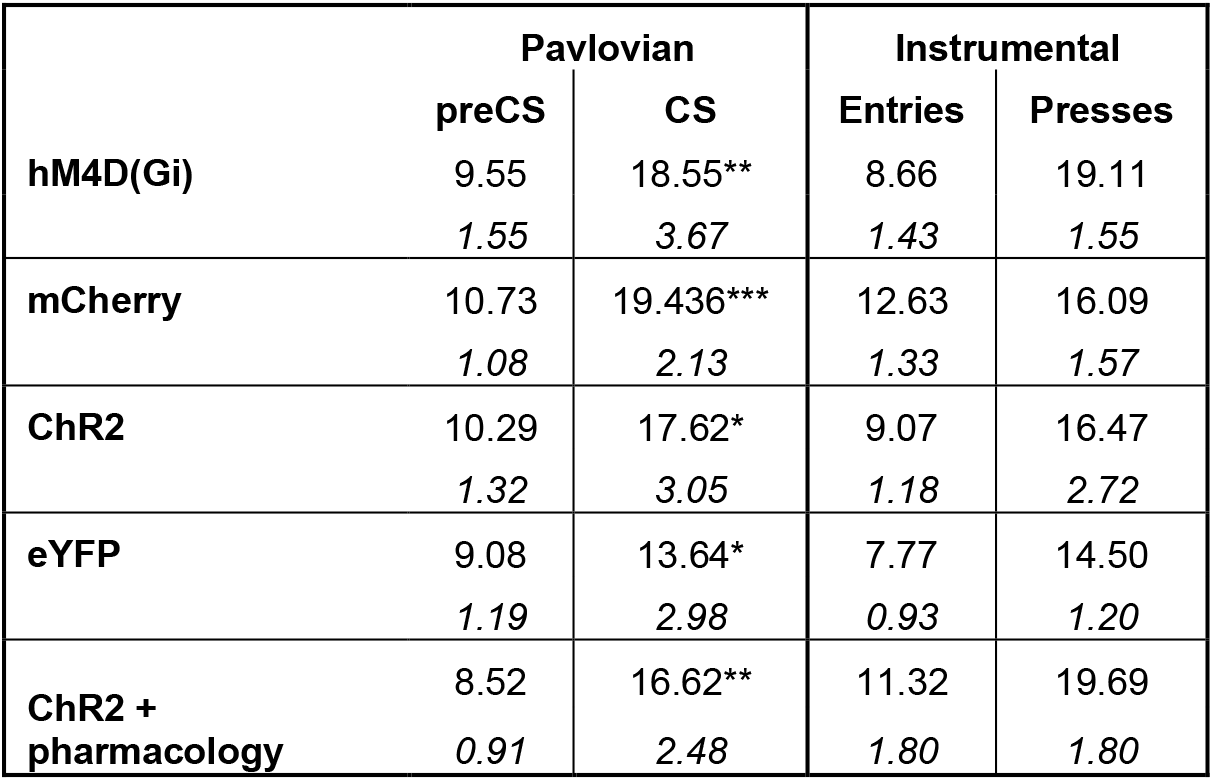
Training data. Training data from the last day of Pavlovian and instrumental training averaged across subjects for each experimental group. Pavlovian data are shown as food-port entries/min (s.e.m. below) averaged across the 2-min baseline periods immediately prior to CS^+^ onset and the food-port entries/min during CS^+^ probe period (after CS onset, but prior to reward delivery). In all cases, rats entered the food port more during the CS probe period than during the baseline periods. Instrumental data are shown as food-port entries/min (Entries) and lever presses/min (Presses) averaged across the entire session. \**P*<0.05, \*\**P*<0.01, \*\*\**P*<0.001.

**Supplemental Figure 1.**
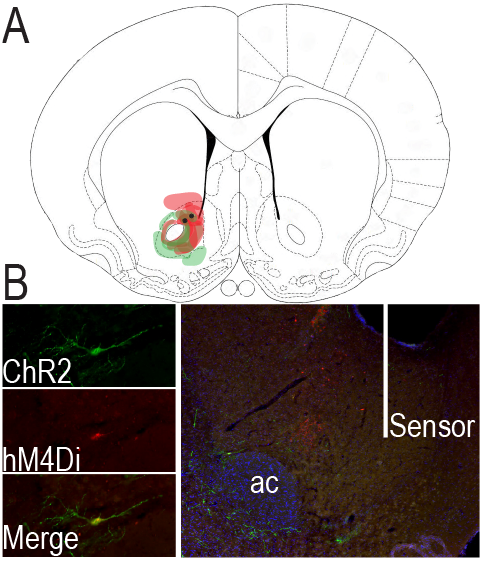
Histological verification of hM4D(Gi) + ChR2 co-expression in NAc cholinergic interneurons. (**A**) Schematic representation of ChR2-eYFP and hM4D(Gi)-mCherry expression and sensor placement the NAc for both subjects. Slide represents 1.00 mm anterior to bregma. (**B**) Representative immunofluorescent image of ChR2-eYFP and hM4D(Gi)-mCherry-expressing cholinergic interneurons in the NAc with biosensor track. AC, anterior commissure.

**Supplemental Figure 2.**
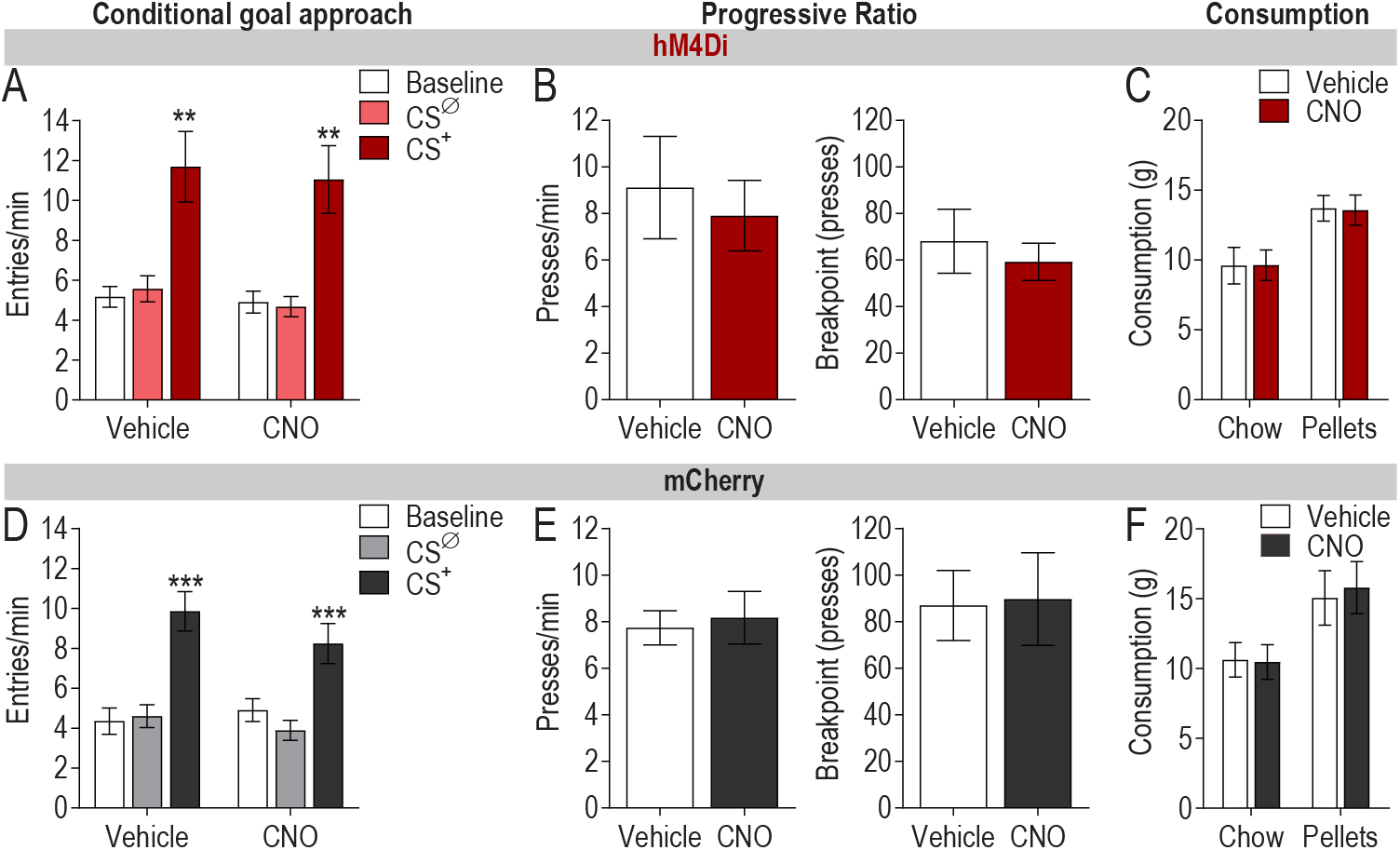
Chemogenetic activation of NAc cholinergic interneurons does not alter conditional approach behavior, lever pressing during a progressive ratio test, or food consumption. (**A**) Effect of chemogenetic inactivation of NAc cholinergic interneurons on Pavlovian conditional food-port approach responding during PIT. *N*=19. Following either vehicle or CNO, the CS^+^ significantly elevated entries into the food-delivery port relative to baseline and CS^Ø^ periods (CS period: *F*_2,36_=19.63, *P*=0.0002; Drug: *F*_1,18_=1.08, *P*=0.31; CS x Drug: *F*_2,36_=0.18, *P*=0.76). (**B**) Following PIT testing, a subset of subjects (*N*=8) were given instrumental retraining then transitioned to a progressive ratio schedule of reinforcement for which after each earned reinforcer the response requirement increased by 50% (rounded up, beginning with a single press requirement). Following 3 days of progressive ratio training, rats were given 2 progressive ratio tests, with intervening retraining, one following vehicle and one following injection of CNO. Inactivation of NAc cholinergic interneurons did not influence either press rate (left; *t*_7_=0.82, *P*=0.44) or break point (right, *t*_7_= 1.14, *P*=0.29) on the progressive ratio. (**C**) Following PIT, rats were tested for the influence of NAc cholinergic interneuron inactivation on consumption of the chocolate pellets (*N*=19) that had been paired with the CS^+^ and lever pressing and of home chow (*N*=8). NAc cholinergic interneuron inactivation with CNO relative to vehicle injection did not influence consumption of either food (Chow: *t*_7_=0.09, *P*=0.93; Pellets: *t*_18_=0.23, *P*=0.82). (**D**) There was no effect of CNO on Pavlovian conditional food-port approach responding during PIT in subjects lacking hM4D(Gi) (CS period: *F*_2,30_=42.97, *P*<0.0001; Drug: *F*_1,15_=0.79, *P*=0.39; CS x Drug: *F*_2,30_=2.51, *P*=0.12) (*N*=16). (**E**) There was no effect of CNO on lever pressing during a progressive ratio test in subjects lacking hM4D(Gi) (Press rate: *t*_8_=0.32, *P*=0.74; Break point: *t*_8_=0.18, *P*=0.86; *N*=9). (**F**) There was no effect of CNO on consumption of either home chow (*t*_8_=0.32, *P*=0.76) or chocolate pellets (*t*_8_=0.68, *P*=0.52) in subjects lacking hM4D(Gi) (*N*=9). Mean +1 s.e.m. \*\*\**P*<0.001.

**Supplemental Figure 3.**
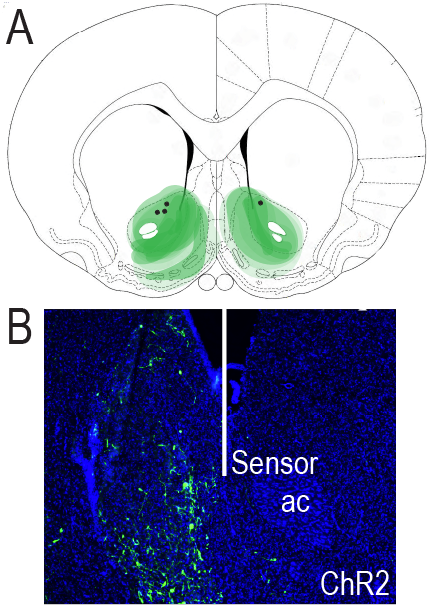
Histological verification of ChR2 expression in NAc cholinergic interneurons for ChR2 validation subjects. (**A**) Schematic representation of ChR2-eYFP expression and sensor placement the NAc for all subjects. Slide represents 0.7 mm anterior to bregma. (**B**) Representative immunofluorescent image of ChR2-eYFP expressing cholinergic interneurons in the NAc with biosensor track. AC, anterior commissure.

**Supplemental Figure 4.**
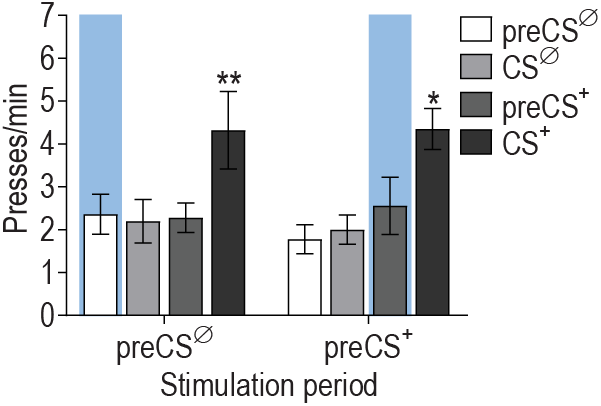
Optical stimulation of NAc cholinergic interneurons has no effect during either preCS^+^ or preCS^Ø^ baseline periods. There were two baseline’ stimulation control periods: one in which NAc cholinergic interneurons were optically stimulated during each of the 2-min periods prior to the CS^+^ and another in which NAc cholinergic interneurons were stimulated during the 2-min preCS^Ø^ periods. In both cases, there was no effect of stimulation on pressing during the baseline period or on the expression of PIT (CS period: *F*_3,24_=9.60, *P*=0.0002; Stimulation period: *F*_1,8_=0.05, *P*=0.82; CS x Stimulation period: *F*_3,24_=0.34, *P*=0.79). Mean +1 s.e.m. \**P*<0.05, \*\**P*<0.01.

**Supplemental Figure 5.**
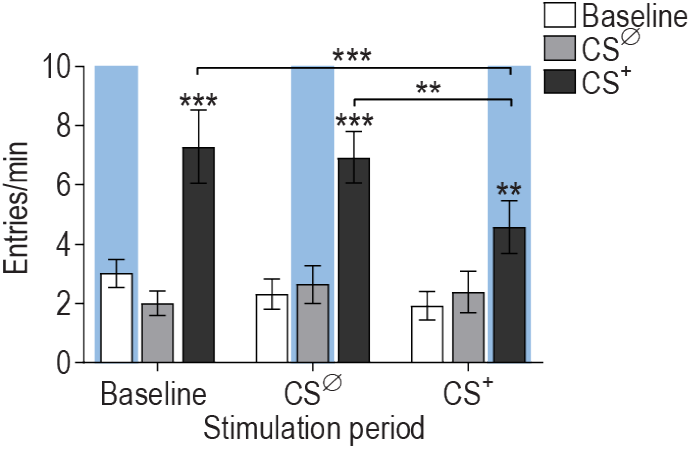
Effect of optical stimulation of NAc cholinergic interneurons on food-port approach responses. Effect of optical stimulation of NAc cholinergic interneurons on food-port approach responding during PIT (CS period: *F*_2,16_=27.9, *P*<0.0001; Stimulation period: *F*_2,16_=2.04, *P*=0.16; CS x Stimulation period: *F*_4,32_=2.979, *P*=0.03; *N*=9). During each test food-port entries were significantly elevated during CS^+^ presentation. Stimulation of NAc cholinergic interneurons concurrent with CS^+^ presentation did cause these to be fewer than during the control baseline or CS^Ø^ stimulation sessions (*P*<0.01). This slight attenuation is expected given that 1) Pavlovian food-port entries are in part sensitive to the same incentive motivational processes supporting PIT and 2) because food-port entries become linked to instrumental actions during training (113, 114). Mean +1 s.e.m. \*\**P*<0.01, \*\*\**P*<0.001. These data indicate that NAc CIN stimulation does not prevent the CS^+^ from eliciting a significant Pavlovian conditional food-port approach response and suggest the subjects are still capable of recognizing the CS^+^.

**Supplemental Figure 6.**
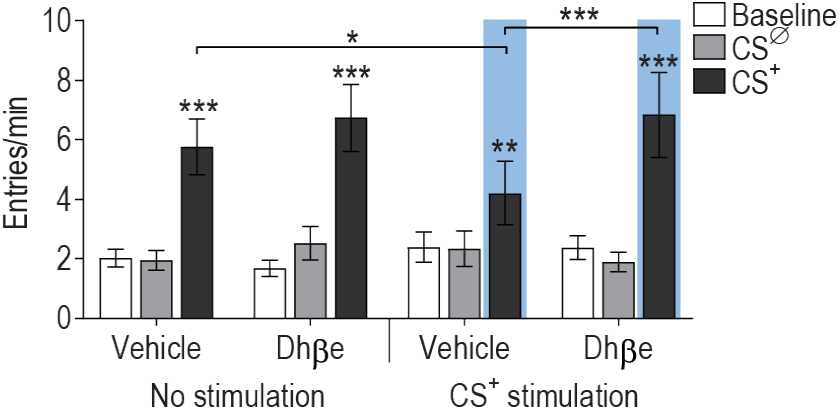
Effect of optical stimulation of NAc cholinergic interneurons and intra-NAc DhβE on food-port approach responses. Effect of optical stimulation of NAc cholinergic interneurons during CS^+^ following either intra-NAc vehicle or DhβE infusion on food-port approach responding during PIT (CS period: *F*_2,20_=22.14, *P*<0.0001; CS^+^ Stimulation: *F*_1,10_=0.10, *P*=0.75; Drug: *F*_1,10_=5.06, *P*=0.048; CS x Drug: *F*_2,20_=8.21, *P*=0.002; Drug x Stimulation: *F*_1,10_=0.08, *P*=0.78; CS x Drug x Stimulation: *F*_2,20_=2.23, *P*=0.13; *N*=11). Food port entries were significantly elevated during the CS^+^ for each test, though stimulation of NAc CINs concurrent with the CS^+^ did slightly attenuate this. Mean +1 s.e.m. \**P*<0.05, \*\**P*<0.01, \*\*\**P*<0.001.

## REFERENCES

1. Corbit LH, Balleine BW (2015): Learning and Motivational Processes Contributing to Pavlovian-Instrumental Transfer and Their Neural Bases: Dopamine and Beyond. Curr Top Behav Neurosci.

2. Cartoni E, Balleine B, Baldassarre G (2016): Appetitive Pavlovian-instrumental Transfer: A review. Neurosci Biobehav Rev. 71:829–848.

3. Johnson AW (2013): Eating beyond metabolic need: how environmental cues influence feeding behavior. Trends Neurosci. 36:101–109.

4. Garbusow M, Schad DJ, Sebold M, Friedel E, Bernhardt N, Koch SP, et al. (2015): Pavlovian-to-instrumental transfer effects in the nucleus accumbens relate to relapse in alcohol dependence. Addict Biol.

5. Garbusow M, Schad DJ, Sommer C, Jünger E, Sebold M, Friedel E, et al. (2014): Pavlovian-to-instrumental transfer in alcohol dependence: a pilot study. Neuropsychobiology. 70:111–121.

6. Corbit LH, Janak PH (2016): Changes in the Influence of Alcohol-Paired Stimuli on Alcohol Seeking across Extended Training. Front Psychiatry. 7:169.

7. Corbit LH, Janak PH (2007): Ethanol-associated cues produce general pavlovian-instrumental transfer. Alcohol Clin Exp Res. 31:766–774.

8. Robinson MJ, Robinson TE, Berridge KC (2013): Incentive salience and the transition to addiction. In: Miller PM, editor. Biological Research on Addiction 2: Academic Press, pp 391–399.

9. Ostlund SB, LeBlanc KH, Kosheleff AR, Wassum KM, Maidment NT (2014): Phasic mesolimbic dopamine signaling encodes the facilitation of incentive motivation produced by repeated cocaine exposure. Neuropsychopharmacology. 39:2441–2449.

10. LeBlanc KH, Ostlund SB, Maidment NT (2012): Pavlovian-to-instrumental transfer in cocaine seeking rats. Behav Neurosci. 126:681–689.

11. Hogarth L, Maynard OM, Munafò MR (2014): Plain cigarette packs do not exert Pavlovian to instrumental transfer of control over tobacco-seeking. Addiction. 110:174–182.

12. Leblanc KH, Maidment NT, Ostlund SB (2013): Repeated cocaine exposure facilitates the expression of incentive motivation and induces habitual control in rats. PLoS One. 8:e61355.

13. Quail SL, Morris RW, Balleine BW (2016): Stress associated changes in Pavlovian-instrumental transfer in humans. Q J Exp Psychol (Hove). 1–29.

14. Morgado P, Silva M, Sousa N, Cerqueira JJ (2012): Stress Transiently Affects Pavlovian-to-Instrumental Transfer. Front Neurosci. 6:93.

15. Pielock SM, Sommer S, Hauber W (2013): Post-training glucocorticoid receptor activation during Pavlovian conditioning reduces Pavlovian-instrumental transfer in rats. Pharmacol Biochem Behav. 104:125–131.

16. Huys QJ, Gölzer M, Friedel E, Heinz A, Cools R, Dayan P, et al. (2016): The specificity of Pavlovian regulation is associated with recovery from depression. Psychol Med. 46:1027–1035.

17. Talmi D, Seymour B, Dayan P, Dolan RJ (2008): Human pavlovian-instrumental transfer. J Neurosci. 28:360–368.

18. Corbit LH, Balleine BW (2011): The general and outcome-specific forms of Pavlovian-instrumental transfer are differentially mediated by the nucleus accumbens core and shell. J Neurosci. 31:11786–11794.

19. Corbit LH, Muir JL, Balleine BW (2001): The role of the nucleus accumbens in instrumental conditioning: Evidence of a functional dissociation between accumbens core and shell. J Neurosci. 21:3251–3260.

20. Scarr E, Gibbons AS, Neo J, Udawela M, Dean B (2013): Cholinergic connectivity: it’s implications for psychiatric disorders. Front Cell Neurosci. 7:55.

21. Dulawa SC, Janowsky DS (2018): Cholinergic regulation of mood: from basic and clinical studies to emerging therapeutics. Mol Psychiatry.

22. Dautan D, Huerta-Ocampo I, Witten IB, Deisseroth K, Bolam JP, Gerdjikov T, et al. (2014): A major external source of cholinergic innervation of the striatum and nucleus accumbens originates in the brainstem. J Neurosci. 34:4509–4518.

23. Zhou FM, Wilson CJ, Dani JA (2002): Cholinergic interneuron characteristics and nicotinic properties in the striatum. J Neurobiol. 53:590–605.

24. Rymar VV, Sasseville R, Luk KC, Sadikot AF (2004): Neurogenesis and stereological morphometry of calretinin-immunoreactive GABAergic interneurons of the neostriatum. J Comp Neurol. 469:325–339.

25. Descarries L, Gisiger V, Steriade M (1997): Diffuse transmission by acetylcholine in the CNS. Prog Neurobiol. 53:603–625.

26. Descarries L, Mechawar N (2000): Ultrastructural evidence for diffuse transmission by monoamine and acetylcholine neurons of the central nervous system. Prog Brain Res. 125:27–47.

27. Wilson CJ, Chang HT, Kitai ST (1990): Firing patterns and synaptic potentials of identified giant aspiny interneurons in the rat neostriatum. J Neurosci. 10:508–519.

28. Inokawa H, Yamada H, Matsumoto N, Muranishi M, Kimura M (2010): Juxtacellular labeling of tonically active neurons and phasically active neurons in the rat striatum. Neuroscience. 168:395–404.

29. Aosaki T, Tsubokawa H, Ishida A, Watanabe K, Graybiel AM, Kimura M (1994): Responses of tonically active neurons in the primate’s striatum undergo systematic changes during behavioral sensorimotor conditioning. J Neurosci. 14:3969–3984.

30. Cachope R, Cheer JF (2014): Local control of striatal dopamine release. Front Behav Neurosci. 8:188.

31. Cragg SJ (2006): Meaningful silences: how dopamine listens to the ACh pause. Trends Neurosci. 29:125–131.

32. Sulzer D, Cragg SJ, Rice ME (2016): Striatal dopamine neurotransmission: regulation of release and uptake. Basal Ganglia.

33. Mark GP, Rada P, Pothos E, Hoebel BG (1992): Effects of feeding and drinking on acetylcholine release in the nucleus accumbens, striatum, and hippocampus of freely behaving rats. J Neurochem. 58:2269–2274.

34. Helm KA, Rada P, Hoebel BG (2003): Cholecystokinin combined with serotonin in the hypothalamus limits accumbens dopamine release while increasing acetylcholine: a possible satiation mechanism. Brain Res. 963:290–297.

35. Warner-Schmidt JL, Schmidt EF, Marshall JJ, Rubin AJ, Arango-Lievano M, Kaplitt MG, et al. (2012): Cholinergic interneurons in the nucleus accumbens regulate depression-like behavior. Proc Natl Acad Sci U S A. 109:11360–11365.

36. Hoebel BG, Avena NM, Rada P (2007): Accumbens dopamine-acetylcholine balance in approach and avoidance. Curr Opin Pharmacol. 7:617–627.

37. Nougaret S, Ravel S (2015): Modulation of Tonically Active Neurons of the Monkey Striatum by Events Carrying Different Force and Reward Information. J Neurosci. 35:15214–15226.

38. Lee IH, Seitz AR, Assad JA (2006): Activity of tonically active neurons in the monkey putamen during initiation and withholding of movement. J Neurophysiol. 95:2391–2403.

39. Ravel S, Legallet E, Apicella P (2003): Responses of tonically active neurons in the monkey striatum discriminate between motivationally opposing stimuli. J Neurosci. 23:8489–8497.

40. Morris G, Arkadir D, Nevet A, Vaadia E, Bergman H (2004): Coincident but distinct messages of midbrain dopamine and striatal tonically active neurons. Neuron. 43:133–143.

41. Joshua M, Adler A, Mitelman R, Vaadia E, Bergman H (2008): Midbrain dopaminergic neurons and striatal cholinergic interneurons encode the difference between reward and aversive events at different epochs of probabilistic classical conditioning trials. J Neurosci. 28:11673–11684.

42. Apicella P, Scarnati E, Schultz W (1991): Tonically discharging neurons of monkey striatum respond to preparatory and rewarding stimuli. Exp Brain Res. 84:672–675.

43. Shimo Y, Hikosaka O (2001): Role of tonically active neurons in primate caudate in reward-oriented saccadic eye movement. J Neurosci. 21:7804–7814.

44. Kimura M, Rajkowski J, Evarts E (1984): Tonically discharging putamen neurons exhibit set-dependent responses. Proc Natl Acad Sci U S A. 81:4998–5001.

45. Apicella P (2007): Leading tonically active neurons of the striatum from reward detection to context recognition. Trends Neurosci. 30:299–306.

46. Aosaki T, Kimura M, Graybiel AM (1995): Temporal and spatial characteristics of tonically active neurons of the primate’s striatum. J Neurophysiol. 73:1234–1252.

47. Bray S, Rangel A, Shimojo S, Balleine B, O’Doherty JP (2008): The neural mechanisms underlying the influence of pavlovian cues on human decision making. J Neurosci. 28:5861–5866.

48. Prévost C, Liljeholm M, Tyszka JM, O’Doherty JP (2012): Neural correlates of specific and general Pavlovian-to-Instrumental Transfer within human amygdalar subregions: a high-resolution fMRI study. J Neurosci. 32:8383–8390.

49. Allman MJ, DeLeon IG, Cataldo MF, Holland PC, Johnson AW (2010): Learning processes affecting human decision making: An assessment of reinforcer-selective Pavlovian-to-instrumental transfer following reinforcer devaluation. J Exp Psychol Anim Behav Process. 36:402–408.

50. Nadler N, Delgado MR, Delamater AR (2011): Pavlovian to instrumental transfer of control in a human learning task. Emotion. 11:1112–1123.

51. Trick L, Hogarth L, Duka T (2011): Prediction and uncertainty in human Pavlovian to instrumental transfer. J Exp Psychol Learn Mem Cogn. 37:757–765.

52. Martinovic J, Jones A, Christiansen P, Rose AK, Hogarth L, Field M (2014): Electrophysiological responses to alcohol cues are not associated with Pavlovian-to-instrumental transfer in social drinkers. PLoS One. 9:e94605.

53. Lovibond PF, Colagiuri B (2013): Facilitation of voluntary goal-directed action by reward cues. Psychological Science. 24:2030–2037.

54. Seabrooke T, Le Pelley ME, Hogarth L, Mitchell CJ (2017): Evidence of a goal-directed process in human Pavlovian-instrumental transfer. J Exp Psychol Anim Learn Cogn. 43:377–387.

55. Lehner R, Balsters JH, Herger A, Hare TA, Wenderoth N (2016): Monetary, Food, and Social Rewards Induce Similar Pavlovian-to-Instrumental Transfer Effects. Front Behav Neurosci. 10:247.

56. Witten IB, Steinberg EE, Lee SY, Davidson TJ, Zalocusky KA, Brodsky M, et al. (2011): Recombinase-driver rat lines: tools, techniques, and optogenetic application to dopamine-mediated reinforcement. Neuron. 72:721–733.

57. Collins AL, Aitken TJ, Greenfield VY, Ostlund SB, Wassum KM (2016): Nucleus Accumbens Acetylcholine Receptors Modulate Dopamine and Motivation. Neuropsychopharmacology.

58. Lichtenberg NT, Pennington ZT, Holley SM, Greenfield VY, Cepeda C, Levine MS, et al. (2017): Basolateral amygdala to orbitofrontal cortex projections enable cue-triggered reward expectations. J Neurosci.

59. Malvaez M, Shieh C, Murphy M, Greenfield V, Wassum K (2018): Distinct cortical-amygdala projections drive reward value encoding and retrieval. BioRxiv.

60. Wassum KM, Ostlund SB, Loewinger GC, Maidment NT (2013): Phasic Mesolimbic Dopamine Release Tracks Reward Seeking During Expression of Pavlovian-to-Instrumental Transfer. Biol Psychiatry. 73:747–755.

61. Wassum KM, Ostlund SB, Balleine BW, Maidment NT (2011): Differential dependence of Pavlovian incentive motivation and instrumental incentive learning processes on dopamine signaling. Learn Mem. 18:475–483.

62. Tabachnick BG, Fidell LS, Osterlind SJ (2001): Using multivariate statistics.

63. Bennett BD, Wilson CJ (1999): Spontaneous activity of neostriatal cholinergic interneurons in vitro. J Neurosci. 19:5586–5596.

64. Wilson CJ, Goldberg JA (2006): Origin of the slow afterhyperpolarization and slow rhythmic bursting in striatal cholinergic interneurons. J Neurophysiol. 95:196–204.

65. Jones IW, Bolam JP, Wonnacott S (2001): Presynaptic localisation of the nicotinic acetylcholine receptor beta2 subunit immunoreactivity in rat nigrostriatal dopaminergic neurones. J Comp Neurol. 439:235–247.

66. Exley R, Cragg SJ (2008): Presynaptic nicotinic receptors: a dynamic and diverse cholinergic filter of striatal dopamine neurotransmission. Br J Pharmacol. 153 Suppl 1:S283–297.

67. Exley R, Clements MA, Hartung H, McIntosh JM, Cragg SJ (2008): Alpha6-containing nicotinic acetylcholine receptors dominate the nicotine control of dopamine neurotransmission in nucleus accumbens. Neuropsychopharmacology. 33:2158–2166.

68. Threlfell S, Lalic T, Platt NJ, Jennings KA, Deisseroth K, Cragg SJ (2012): Striatal dopamine release is triggered by synchronized activity in cholinergic interneurons. Neuron. 75:58–64.

69. Cachope R, Mateo Y, Mathur BN, Irving J, Wang HL, Morales M, et al. (2012): Selective activation of cholinergic interneurons enhances accumbal phasic dopamine release: setting the tone for reward processing. Cell Rep. 2:33–41.

70. Rice ME, Cragg SJ (2004): Nicotine amplifies reward-related dopamine signals in striatum. Nat Neurosci. 7:583–584.

71. Zhou FM, Liang Y, Dani JA (2001): Endogenous nicotinic cholinergic activity regulates dopamine release in the striatum. Nat Neurosci. 4:1224–1229.

72. Berridge KC (2007): The debate over dopamine’s role in reward: the case for incentive salience. Psychopharmacology (Berl). 191:391–431.

73. Aitken TJ, Greenfield VY, Wassum KM (2016): Nucleus accumbens core dopamine signaling tracks the need-based motivational value of food-paired cues. J Neurochem. 136:1026–1036.

74. Saunders BT, Richard JM, Margolis EB, Janak PH (2018): Dopamine neurons create Pavlovian conditioned stimuli with circuit-defined motivational properties. Nat Neurosci. 21:1072–1083.

75. Lex A, Hauber W (2008): Dopamine D1 and D2 receptors in the nucleus accumbens core and shell mediate Pavlovian-instrumental transfer. Learn Mem. 15:483–491.

76. Collins AL, Greenfield VY, Bye JK, Linker KE, Wang AS, Wassum KM (2016): Dynamic mesolimbic dopamine signaling during action sequence learning and expectation violation. Sci Rep. 6:20231.

77. Saunders BT, Robinson TE (2012): The role of dopamine in the accumbens core in the expression of Pavlovian-conditioned responses. Eur J Neurosci. 36:2521–2532.

78. Peciña S, Berridge KC (2013): Dopamine or opioid stimulation of nucleus accumbens similarly amplify cue-triggered ‘wanting’ for reward: entire core and medial shell mapped as substrates for PIT enhancement. Eur J Neurosci.

79. Wyvell CL, Berridge KC (2000): Intra-accumbens amphetamine increases the conditioned incentive salience of sucrose reward: enhancement of reward “wanting” without enhanced “liking” or response reinforcement. J Neurosci. 20:8122–8130.

80. Zhang L, Doyon WM, Clark JJ, Phillips PE, Dani JA (2009): Controls of tonic and phasic dopamine transmission in the dorsal and ventral striatum. Mol Pharmacol. 76:396–404.

81. Zhang H, Sulzer D (2004): Frequency-dependent modulation of dopamine release by nicotine. Nat Neurosci. 7:581–582.

82. Threlfell S, Cragg SJ (2011): Dopamine signaling in dorsal versus ventral striatum: the dynamic role of cholinergic interneurons. Front Syst Neurosci. 5:11.

83. Lim SA, Kang UJ, McGehee DS (2014): Striatal cholinergic interneuron regulation and circuit effects. Front Synaptic Neurosci. 6:22.

84. Rice ME, Cragg SJ (2004): Nicotine amplifies reward-related dopamine signals in striatum. Nat Neurosci. 7:583–584.

85. Stouffer MA, Woods CA, Patel JC, Lee CR, Witkovsky P, Bao L, et al. (2015): Insulin enhances striatal dopamine release by activating cholinergic interneurons and thereby signals reward. Nat Commun. 6:8543.

86. Koob GF (1999): Corticotropin-releasing factor, norepinephrine, and stress. Biol Psychiatry. 46:1167–1180.

87. Koob GF, Bloom FE (1985): Corticotropin-releasing factor and behavior. Fed Proc. 44:259–263.

88. Lemos JC, Wanat MJ, Smith JS, Reyes BA, Hollon NG, Van Bockstaele EJ, et al. (2012): Severe stress switches CRF action in the nucleus accumbens from appetitive to aversive. Nature. 490:402–406.

89. Koob GF, Heinrichs SC, Pich EM, Menzaghi F, Baldwin H, Miczek K, et al. (1993): The role of corticotropin-releasing factor in behavioural responses to stress. Ciba Found Symp. 172:277–289; discussion 290-275.

90. Lemos J, Shin J, Ingebretson A, Dobbs L, Alvarez V (2018): Cholinergic interneurons as a novel target of CRF in the striatum that is spared by repeated stress. bioRxiv.

91. Chen YW, Rada PV, Bützler BP, Leibowitz SF, Hoebel BG (2012): Corticotropin-releasing factor in the nucleus accumbens shell induces swim depression, anxiety, and anhedonia along with changes in local dopamine/acetylcholine balance. Neuroscience. 206:155–166.

92. Dölen G, Darvishzadeh A, Huang KW, Malenka RC (2013): Social reward requires coordinated activity of nucleus accumbens oxytocin and serotonin. Nature. 501:179–184.

93. Virk MS, Sagi Y, Medrihan L, Leung J, Kaplitt MG, Greengard P (2016): Opposing roles for serotonin in cholinergic neurons of the ventral and dorsal striatum. Proc Natl Acad Sci U S A. 113:734–739.

94. Olney JJ, Warlow SM, Naffziger EE, Berridge KC (2018): Current perspectives on incentive salience and applications to clinical disorders. Curr Opin Behav Sci. 22:59–69.

95. Robinson TE, Berridge KC (1993): The neural basis of drug craving: an incentive-sensitization theory of addiction. Brain Res Brain Res Rev. 18:247–291.

96. Hikida T, Kitabatake Y, Pastan I, Nakanishi S (2003): Acetylcholine enhancement in the nucleus accumbens prevents addictive behaviors of cocaine and morphine. Proc Natl Acad Sci U S A. 100:6169–6173.

97. Hikida T, Kaneko S, Isobe T, Kitabatake Y, Watanabe D, Pastan I, et al. (2001): Increased sensitivity to cocaine by cholinergic cell ablation in nucleus accumbens. Proc Natl Acad Sci U S A. 98:13351–13354.

98. Treadway MT, Zald DH (2013): Parsing Anhedonia: Translational Models of Reward-Processing Deficits in Psychopathology. Curr Dir Psychol Sci. 22:244–249.

99. Paxinos G, Watson C (1998): The rat brain in stereotaxic coordinates. 4th ed.: Academic Press.

100. Stefanik MT, Kalivas PW (2013): Optogenetic dissection of basolateral amygdala projections during cue-induced reinstatement of cocaine seeking. Front Behav Neurosci. 7:213.

101. Malvaez M, Greenfield VY, Wang AS, Yorita AM, Feng L, Linker KE, et al. (2015): Basolateral amygdala rapid glutamate release encodes an outcome-specific representation vital for reward-predictive cues to selectively invigorate reward-seeking actions. Sci Rep. 5:12511.

102. Smith KS, Bucci DJ, Luikart BW, Mahler SV (2016): DREADDS: Use and application in behavioral neuroscience. Behav Neurosci. 130:137–155.

103. Wassum KM, Ostlund SB, Maidment NT, Balleine BW (2009): Distinct opioid circuits determine the palatability and the desirability of rewarding events. Proc Natl Acad Sci U S A. 106:12512–12517.

104. Corrigall WA, Coen KM, Adamson KL (1994): Self-administered nicotine activates the mesolimbic dopamine system through the ventral tegmental area. Brain Res. 653:278–284.

105. Wassum KM, Tolosa VM, J. W, Walker E, Monbouquette HG, Maidment NT (2008): Silicon Wafer-Based Platinum Microelectrode Array Biosensor for Near Real-Time Measurement of Glutamate In Vivo. Sensors. 8:5023–5036.

106. Wassum KM, Tolosa VM, Tseng TC, Balleine BW, Monbouquette HG, Maidment NT (2012): Transient Extracellular Glutamate Events in the Basolateral Amygdala Track Reward-Seeking Actions. J Neurosci. 32:2734–2746.

107. Burmeister JJ, Moxon K, Gerhardt GA (2000): Ceramic-based multisite microelectrodes for electrochemical recordings. Anal Chem. 72:187–192.

108. Giuliano C, Parikh V, Ward JR, Chiamulera C, Sarter M (2008): Increases in cholinergic neurotransmission measured by using choline-sensitive microelectrodes: enhanced detection by hydrolysis of acetylcholine on recording sites? Neurochem Int. 52:1343–1350.

109. Parikh V, Pomerleau F, Huettl P, Gerhardt GA, Sarter M, Bruno JP (2004): Rapid assessment of in vivo cholinergic transmission by amperometric detection of changes in extracellular choline levels. Eur J Neurosci. 20:1545–1554.

110. Parikh V, Sarter M (2006): Cortical choline transporter function measured in vivo using choline-sensitive microelectrodes: clearance of endogenous and exogenous choline and effects of removal of cholinergic terminals. Journal of neurochemistry. 97:488–503.

111. Wassum KM, Tolosa VM, Wang J, Walker E, Monbouquette HG, Maidment NT (2008): Silicon Wafer-Based Platinum Microelectrode Array Biosensor for Near Real-Time Measurement of Glutamate in Vivo. Sensors (Basel). 8:5023–5036.

112. Hersman S, Cushman J, Lemelson N, Wassum K, Lotfipour S, Fanselow MS (2017): Optogenetic excitation of cholinergic inputs to hippocampus primes future contextual fear associations. Sci Rep. 7:2333.

113. Marshall AT, Ostlund SB (2018): Repeated cocaine exposure dysregulates cognitive control over cue-evoked reward-seeking behavior during Pavlovian-to-instrumental transfer. Learn Mem. 25:399–409.

114. Halbout B, Marshall A, Azimi A, Liljeholm M, Mahler S, Wassum K, et al. Ventral tegmental dopamine inputs to the nucleus accumbens mediates cue-triggered motivation but not reward expectancy. BioRxiv.

